# BiSEK: a platform for a reliable differential expression analysis

**DOI:** 10.1101/2021.02.22.432271

**Authors:** Roni Haas, Dean Light, Yahav Festinger, Neta Friedman, Ayelet T. Lamm

## Abstract

Differential Expression Analysis (DEA) of RNA-sequencing data is frequently performed for detecting key genes, affected across different conditions. Although DEA-workflows are well established, preceding reliability-testing of the input material, which is crucial for consistent and strong results, is challenging and less straightforward. Here we present Biological Sequence Expression Kit (BiSEK), a graphical user interface-based platform for DEA, dedicated to a reliable inquiry. BiSEK is based on a novel algorithm to track discrepancies between the data and the statistical model design. Moreover, BiSEK enables differential-expression analysis of groups of genes, to identify affected pathways, without relying on the significance of genes comprising them. Using BiSEK, we were able to improve previously conducted analysis, aimed to detect genes affected by FUBP1 depletion in chronic myeloid leukemia cells of mice bone-marrow. We found affected genes that are related to the regulation of apoptosis, supporting *in-vivo* experimental findings. We further tested the host response following SARS-CoV-2 infection. We identified a substantial interferon-I reaction and low expression levels of TLR3, an inducer of interferon-III (IFN-III) production, upon infection with SARS-CoV-2 compared to other respiratory viruses. This finding may explain the low IFN-III response upon SARS-CoV-2 infection. BiSEK is open-sourced, available as a web-interface.

## INTRODUCTION

In genomics studies, a common scientific interest is evaluating quantitative differences across samples, usually based on High-Throughput Sequencing (HTS) data, by conducting Differential Expression Analysis (DEA). DEA is most commonly applied to RNA-sequencing (RNA-seq) datasets, to evaluate gene expression levels of an organism (1). However, DEA is not confined to gene expression, and can be used for other quantitative assessments, such as the measurement of tax abundance differences in Metagenomics, or the assessment of variation in DNA methylation levels. For DEA based on RNA-seq, a typical pre-processing pipeline includes: 1. Mapping of RNA-seq reads to the genome or the transcriptome 2. Counting the number of reads aligned to each gene/transcript/exon for every tested sample to evaluate the expression levels 3. Summarizing the read counts of all samples into a table of counts. Following pre-processing, the count tables are statistically analyzed using DEA models, to infer which genes are significantly altered between samples. Reliable DEA often requires different methods for normalization of read counts, calculation of log-fold-changes and ranking of Differentially Expressed Genes (DEG) by p-values (2).

Various DEA methodologies have been developed in the last decade, to answer the growing demand for practical guidance and the need for statistical methods in detecting quantitative changes in biological experiments. Some of the DEA methodologies are available as software, among them are baySeq (3), DESeq (4), edgeR (5), EBSeq (6), SAMseq (7), DESeq2 (8), limma (9), NOIseq (10) and Sleuth (11).

DESeq is an R/Bioconductor package presenting a DEA methodology based on negative binomial distribution to model read counts, and linking of variance and mean by local regression (4). In 2014, DESeq2, considered as the successor of DESeq, came to light (8). DESeq2 became a highly cited tool (12) and was found to be one of the most sensitive and precise software that exists for DEA (1). DEseq2 suggests advanced extensions and novel features, foremost among them is the use of Empirical Bayes shrinkage-estimators for log fold change (LFC) values. LFC shrinkage is a good solution to deal with low gene count values, which often show small signal to noise ratio, that leads to an improved gene ranking (8). However, the usage of DESeq2 may be challenging for many researchers because it demands computational skills, and particularly programming proficiency in R language. This holds true particularly when analyzing complex study designs deviating from the standard workflow due to multiple conditions, interactions and specialized research questions. For non-programmers, enabling the tool usage through a friendly interface, for which programming skills are not needed, is invaluable.

Several applications were built to ease DESeq2 usage through a Graphical User Interface (GUI), among them are DEApp (13), Shaman (14), and UTAP (15). The mentioned tools enable a DEA analysis given a gene count table (some of the tools enable upstream pre-processing analysis), including the selection of conditions in either single or multiple factor designs, filtration according to a minimum gene count threshold, DESeq2 run and several data visualization options. However, these applications mostly restrict the user to the basic DESeq2 analysis using *results* function (16), and disallow controlling important options that may extensively affect the results and control their correctness. In addition, and more importantly, using these tools or others (whether designed with a user interface or not) does not guarantee the correctness and the accuracy of the results, since these tools do not provide sufficient guiding in a process of verification of the fitness of the input data prior to DEA. Reliable data analysis requires examining the data behavior and verifying the fit of the data to the complexity of the inferences desired by the user.

For a preliminary exploration of the data behavior, it is recommended to perform a Principal Component Analysis (PCA). The PCA reflects the gene-expression variation, under a dimensional reduction that aims to maximally preserve the data variance, by transforming the count data into orthogonal primary components that show the highest variance between samples (17). DESeq2 provides its own PCA implementation which incorporates specific normalization steps that account for the biological nature of the data. The applications mentioned above, support performing the PCA through a GUI. However, while performing PCA is invaluable for correctly analyzing the data, it requires mathematical and statistical knowledge and provides another, possibly more challenging, hurdle for researchers to clear. Incorrect interpretation of the PCA may in some cases lead to serious errors. Often the consistency of biological replicates is not satisfying and the desired variation between different conditions is biased due to discrepancies in the data (i.e. outlier samples, samples that were accidently switched). Therefore, identification of discrepancies in the data and in the relation between the data and the statistical model design is crucial to display reliable and strong DEA results.

To fill the gap of lacking robust DEA tools to examine the data variation in relation to the experimental design, and at the same time to enable seamless DEA of complex studies, we developed Biological Sequence Expression Kit (BiSEK), an automated tool oriented for a reliable and accurate DEA, which can be applied to any quantitative data (not restricted to gene expression). Firstly, we introduced a novel algorithm, PaDETO (Partition Distance Explanation Tree Optimizer), that tracks discrepancies in the data, alerts about problems and offers the best solutions considering the user setup, to increase reliability of the DEA output. Secondly, BiSEK allows adjusting more DESeq2 parameters than other DESeq2 interfaces, enabling non-programmers to better fit statistical models to their scientific question. Thirdly, BiSEK provides a module for differential expression of groups of genes, while DESeq2 only provides differential expression of individual genes. BiSEK is an open-source tool that can be easily locally installed, or be accessed via a web interface.

To demonstrate the power of BiSEK, we applied it to previously published RNA-seq data. We show that BiSEK was able to detect inconsistency in the data, resulting in improved strength of the test to detect affected genes. Analyzing data from FUBP1 depletion in Chronic Myeloid Leukemia (CML) cells of mice bone marrow that was generated by (18), we increased the power of the test and suggested a list of genes responsible for apoptosis, observed directly only experimentally in the study (18). We further tested the host response following SARS-CoV-2 infection, compared to that caused by other respiratory viruses (19). Using BiSEK we were able to show a substantial IFN-I response following SARS-CoV-2 infection that was underestimated by others, using different analysis techniques. Moreover, we suggest that a specific gene, TLR3, which is related to interferon-III (IFN-III) production regulation, rather than an entire process is responsible for the low IFN-III response upon SARS-CoV-2 infection.

## MATERIAL AND METHODS

### BiSEK implementation

BiSEK is written mainly in python (20), with calls to R (21) via the rpy2 library. R is used to run the DESeq2(8) package which is the backbone of the BiSEK App, and the HCPC (22) function in the RFactoMine package which is used as a subroutine in the PaDETO algorithm. The frontend is implemented using Plotly’s Dash library (23). The backend and the PaDETO algorithm use the Numpy (24), Pandas, Dask (25), scikit-learn (26), networkX (27) and matplotlib (28) libraries in python. The app is served using Docker containers (29).

### BiSEK usage of different DESeq2 options

DESeq2 package in R enables applying different methods. To keep this advantage in BiSEK we allow a wide array of analysis options in a graphical user interface (**Figure 1)**:

**Figure 1:**
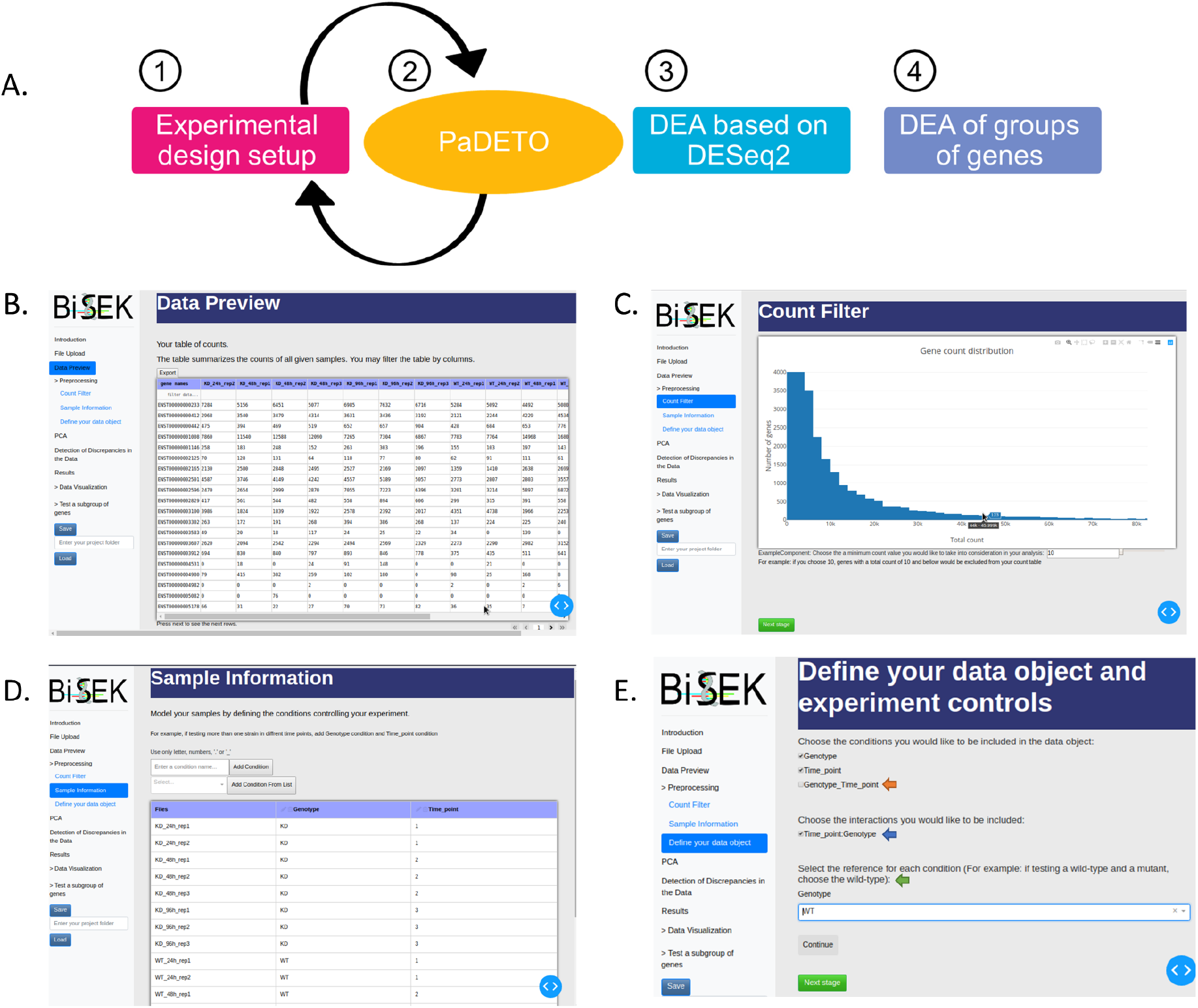
BiSEK preprocessing and experimental design setup workflow demonstration. **(A)** BiSEK’s schematic view. **(B)** Gene-count table after upload. **(C)** Visualization of gene-count distribution, updated in real time during filtration by minimal counts threshold. **(D)** Definition of conditions. **(E)** Designing the data object and reference level definition. Conditions defined by the user and the conditions created by the software are displayed. Light blue scroller, marks the currant stage in BiSEK application. Orange arrow, combination of the defined conditions into a single factor. Blue arrow, interaction term. Green arrow, sets the factor level which the user wants to compare against.

#### Count filtering

We enable filtration the input count table by the sum of all samples counts for a given gene. The default minimal counts threshold is 10 (**Figure 1C**).

#### Defining the data object

One or more of the conditions defined in the sample information table must be chosen to be included in the data object. We suggest adding interaction-terms into the design, and combine the defined factors into a single factor to create a single condition (**Figure 1E**). By default, all conditions defined in the sample information page would be included in the design, but not interaction-terms or combined factors.

#### DESeq2 result table

We allow choosing to apply shrinkage of LFC values using *lfcShrink* function, while both coef and contrast methods are available. Dictated by DESeq2, using coef, but not contrast, enables using the apeglm shrinkage estimator type which is the most recommended by DESeq2 (**Figure S1A, purple arrow)**. However, using contrast permits comparing between several groups within a certain factor while coef restricts the user to use only the indexes of the *resultsNames* DESeq2 function (16) for the test. We allow selecting between coef and contrast methods to enable an informed choice based on one’s needs. If the coef method was selected, we enable setting the type of shrinkage estimator to be normal or apeglm. When no shrinkage is desired by the user we use the DESeq2 *results* function, and permits defining a significance level, alpha, to set a *p* value cutoff, and pAdjustMethod to choose the preferable method for adjusting p-value (**Figure S1B)**. The default setting is dictated by DESq2.

#### lfcThreshold option

A common practice among researchers is to filter out genes with LFC threshold, which is below a certain value, after conducting a statistical significance test according to conventional null hypothesis (LFC=0), to track DEG. However, if the genes of interest are only those with LFC that is below a certain threshold, it is preferable to evaluate it statistically directly, namely defining it in filtering is defined by the user. Further steps may include one or both visualization and Welch two-sample T-test (see below).

#### Visualization

The average normalized counts of the tested samples are plotted against each other on log scale plots. Genes can be colored according to their group belongness, regression lines may be displayed, and gene data labels optionally printed.

#### Welch two-sample T-test

the user may select 2 of the groups to perform a Welch two-sample T-test. In the test, two populations are defined according to the user selection. The null hypothesis is that the Log2FoldChange means of the two groups are equal. Namely, if the null hypothesis is falsified, a significant difference between the expression of a given subgroup of genes and a second group of genes (all genes or another subgroup of genes) is observed. Using the tool, the mean Log2foldchange values of the two groups, the T-test result and a dynamic plot as described in Visualization can be exported.

For all analysis performed using this module in the current study, for either visualization or Welch two-sample T-test, lists were filtered according to their Coefficient Variation (CV) (CV<=1) unless specified.

### PaDETO algorithm

PaDETO takes the following steps: 1) Cluster the PCA data of the experiment (**Figure 2A**). 2) Calculate the distance between the PCA clustering partition of the samples and the experimental setup partition of the samples (**Figure 2B**). 3) If the distance is not zero, look for an explanation that explains the difference between the partitions (**Figure 2C-D**). 4) Return the best series of explanations found (**Figure 2E**). A detailed description of the PaDETO algorithm is in the Supplementary Material, and a schematic view is illustrated in **Figure 2**.

**Figure 2:**
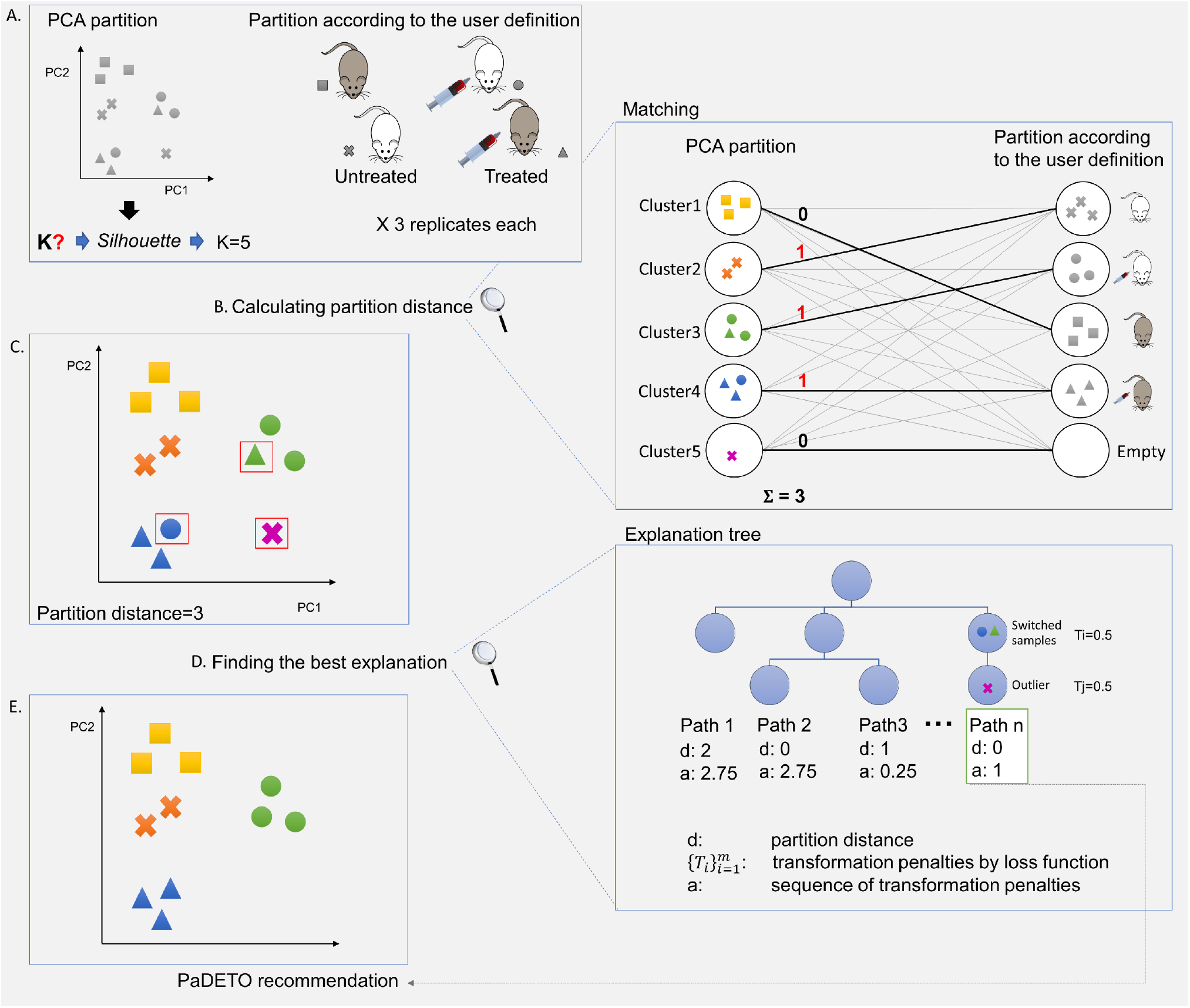
Schematic outline of the PaDETO algorithm. **(A)** Data partition is received by clustering the samples based on the PCA vectors (Left). In this case, 12 samples are partitioned into 5 clusters. A user partition of the data is derived from the sample metadata that the user provides (Right). Here, there are four type of samples with 3 replicas each, classified according to two conditions (Genotype: white or grey mice; Treatment: treated or untreated), each denoted with a unique shape (square: grey, untreated; cross: white, untreated; circle: white, treated; triangular: grey, treated). (**B)** The difference between the Data partition and the user partition is computed using a bipartite graph matching algorithm, with the data partition on the left and the user partition on the right. In this case the partition distance is 3 since at minimum 3 different evaluations would lead to a perfect match. Namely, a perfect match would be achieved if moving: the purple cross to cluster 2, the green triangle to cluster 4, and the blue circle to cluster 3. **(C)** Given the partition, PaDETO knows which samples do not fit the experimental setup. **(D)** Finding the best explanation by graph optimization techniques. Each blue node in the directed graph is an experiment, with the root being the original experiment. Each edge is an explanation, a transformation of the experiment according to a pre-supplied set of sensible transformations (e.g. switching samples, removing an outlier and etc.) each with a given weight. PaDETO considers the shortest weighted path between the root and the best experiments in the graph, according to the partition distance. In this case, switching the blue circle and the green triangle, and then removing the purple cross as an outlier, leads to d=0 while accruing a weight of 1. (**E)** PaDETO presents the modified data according to the chosen paths and supplies a recommendation to the user. The recommendation would follow the selected explanation path. The explanation text is omitted for brevity in this figure. A detailed description of the PaDETO algorithm is given in the Supplementary Material.

### Datasets used in this study and BiSEK’s analysis setup

For datasets used in this study, raw gene counts, were either created by us: (30), GEO accession number GSE122015, **Figure 3**, or downloaded from the GEO: (A) **Figure S2**, GEO accession number GSE140066 (B) **Figure 4**, GEO accession number GSE124304 (18) (C) **Figure 5-6** and **Figures S3-S11**, GEO accession number GSE147507 (19). The results for all parts were created by following the GUI flow with BiSEK default parameters, unless specified.

**Figure 3:**
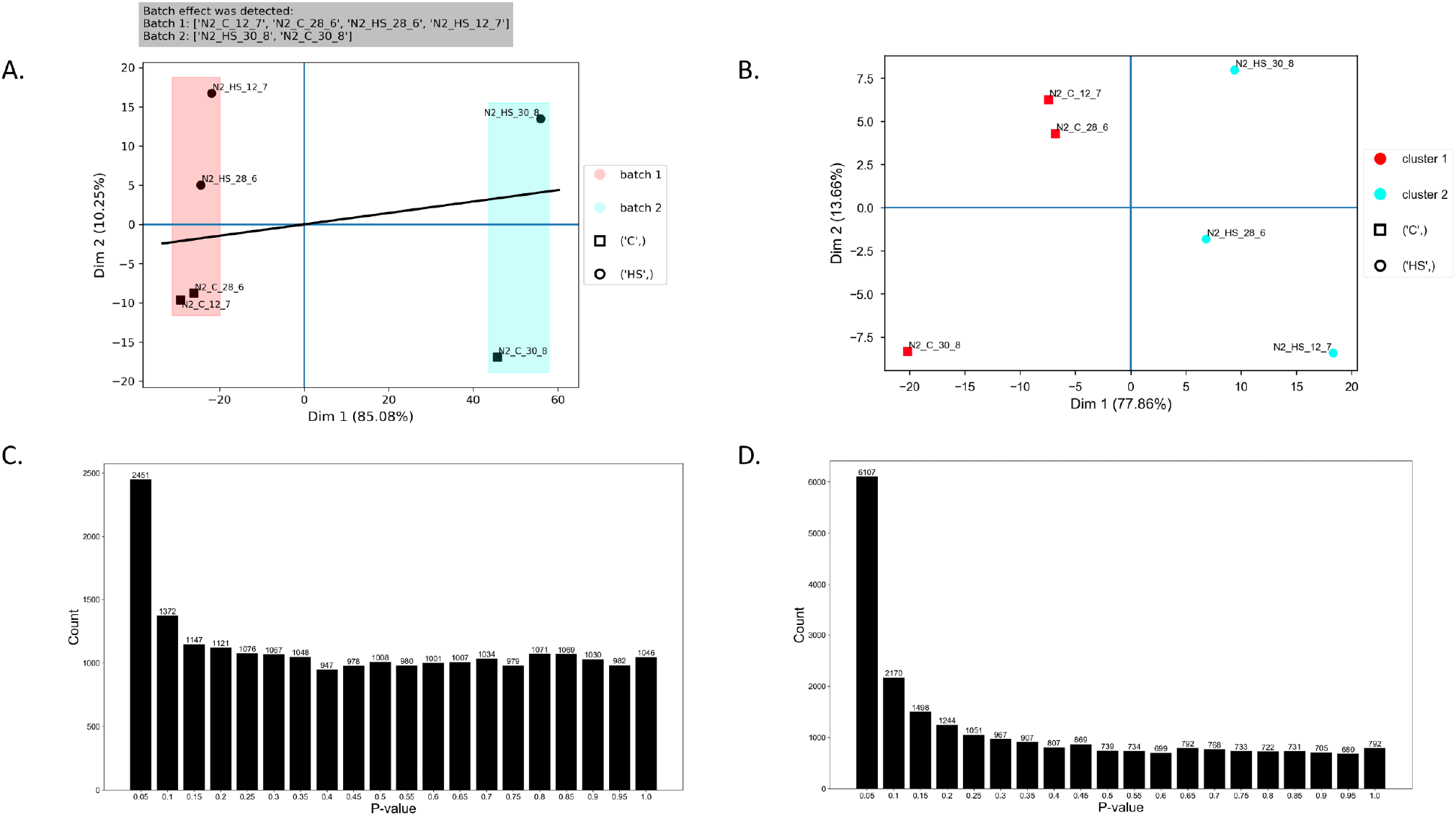
Batch effect detection by PaDETO algorithm leads to improved power of the statistical test. (A) PaDETO identified a batch effect. Colors represent different batches. Shapes represent the expected sample partition as defined by the user. The black line is the linear separator by which the batch effect was defined (see Supplementary Material). (B) The PaDETO partition following batch effect variation removing. The figure is automatically presented by the software when a batch effect is detected. Colors represent the clusters by which PaDETO partitioned the samples based on their variation. Shapes represent the expected partition as defined by the user. The PCA data fits the metadata, as all circle shapes samples are in one cluster (light blue) and all square shaped samples are in another cluster (red) (C-D) P-value distribution following DESeq2 run. (C) Before batch effect is defined. (D) Batch effect condition was added to the sample information table according to PaDETO recommendation. The power of the test in D is higher than in C, indicated by the much larger excess of P-values in the lowest bins (P ≤ 0.10) and correspondingly, much less P-values across higher bins in D, compared to C.

**Figure 4:**
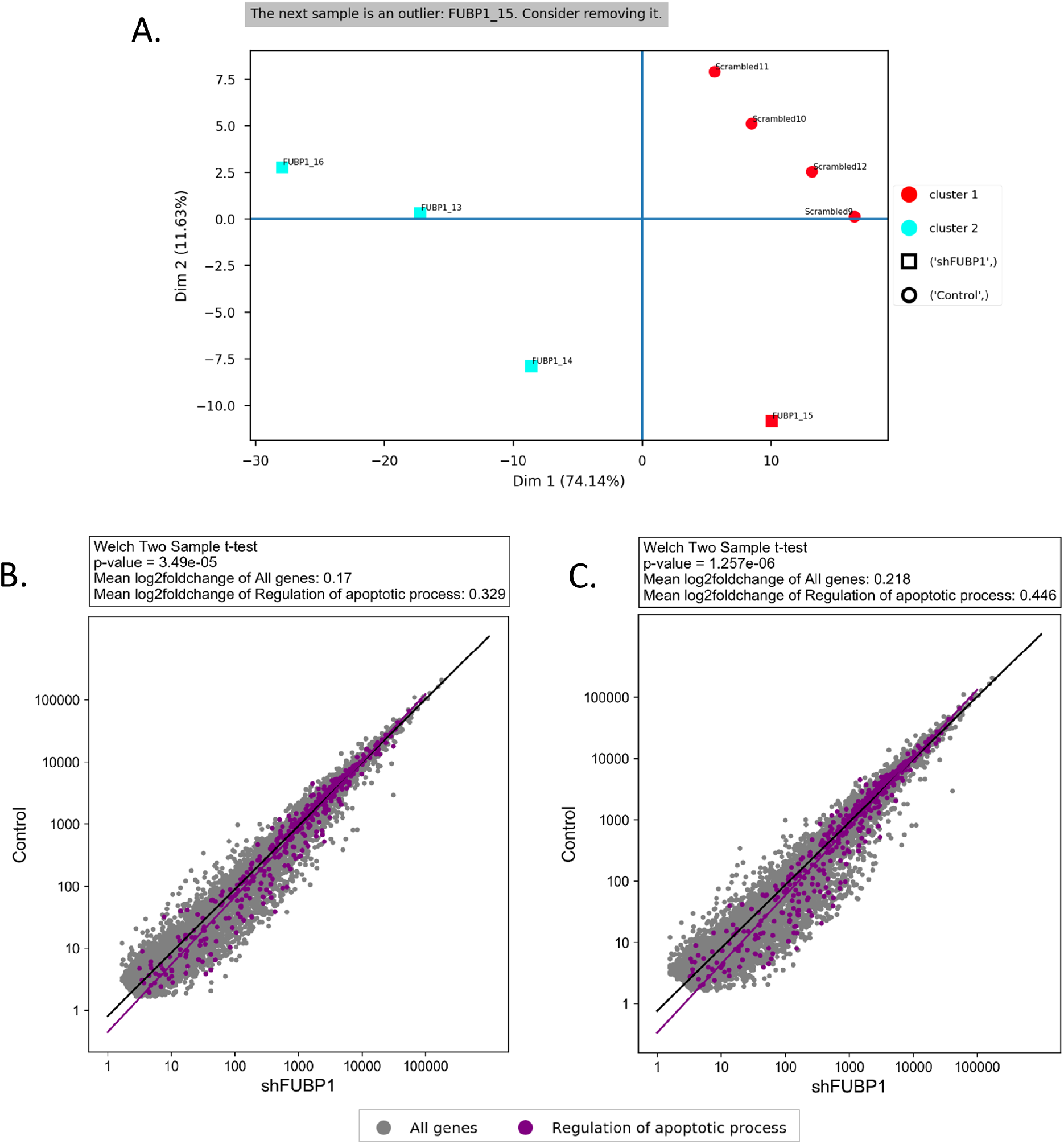
Outlier detection by PaDETO leads for identifying highly influenced genes, by analyzing groups of genes using BiSEK’s module. **(A)** PaDETO’s PCA interpretation pointing to one of the FUBP1 shRNA+ samples as an outlier. Colors represent the clusters by which PaDETO partitioned the samples based on their variation. Shapes represent the expected partition as defined by the user. It could be noticed that FUBP1_15 sample was attributed to the red cluster together with the control scrambled shRNA+ samples (circle shaped), rather than the light blue cluster that includes all other FUBP1 shRNA+ samples (square shaped). **(B-C)** A module for testing deferential expression analysis of groups of genes showing that genes participating in “The regulation of apoptotic process (GO:0042981)” (purple) are significantly upregulated, compared to all genes (grey), when including all samples in the analysis as originally done **(B)** and when excluding the outlier sample **(C)**. Omitting the outlier sample results with a more significant result (P-values are 1.257e-06 in **C**, compared to 3.490e-05 in **B**). Black line is the regression line for all genes. Purple line is the regression line of the apoptotic genes.

**Figure 5:**
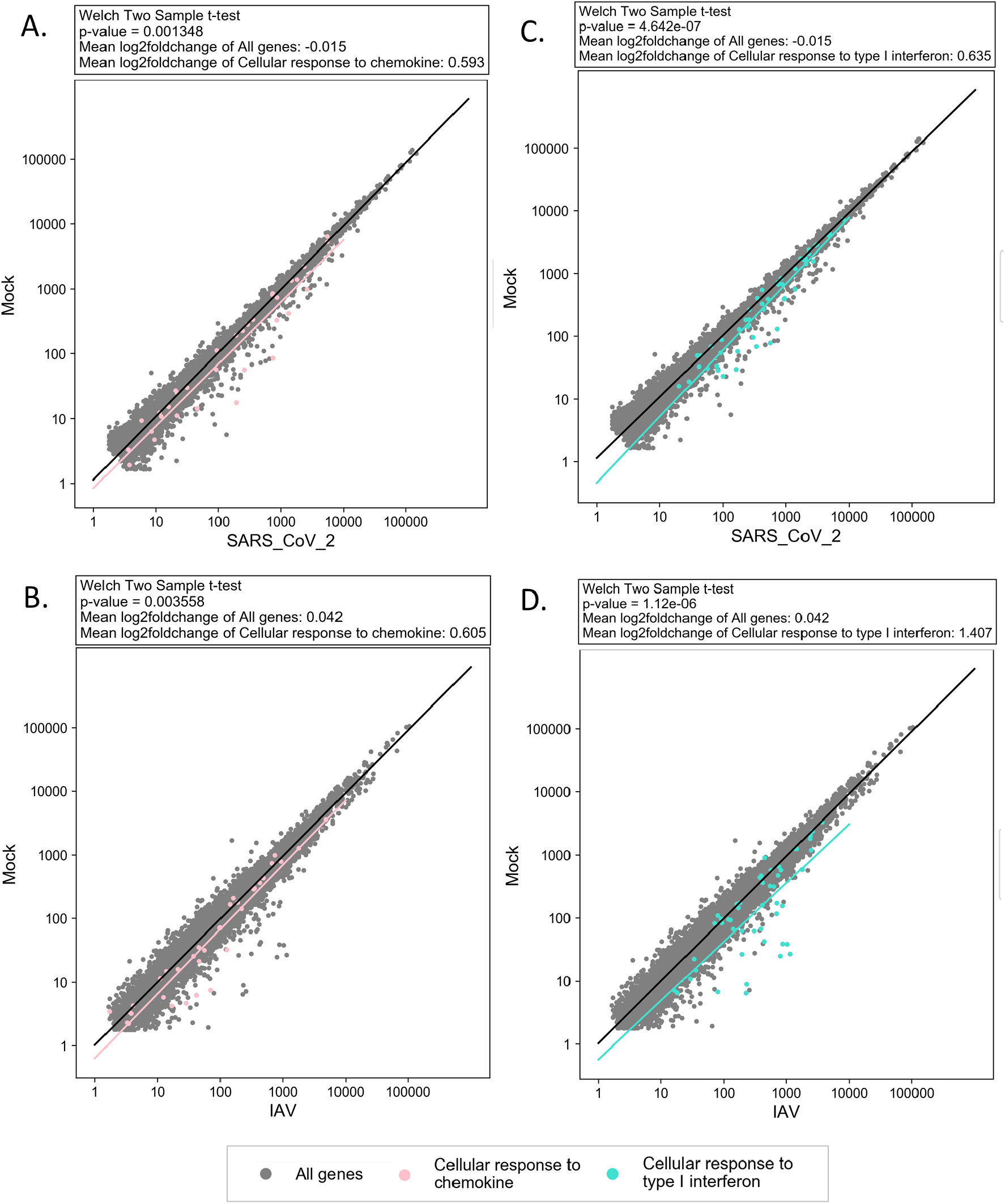
Strong chemokine and IFN-I cell response following SARS-CoV-2 and IAV infection for NHBE cells. Genes related to chemokine activity (pink) are significantly upregulated, compared to all genes (grey) upon SARS-CoV-2 (**A**) and IAV (**B**) infection. Genes related to IFN-I activity (turquoise) are significantly upregulated, compared to all genes (grey) upon SARS-CoV-2 (**C**) and IAV (**D**) infection. The black, pink, and turquoise lines are the regression lines for: all genes, chemokine related genes, and IFN-I related genes, respectively. Log scale plots present average normalized gene counts.

**Figure 6.**
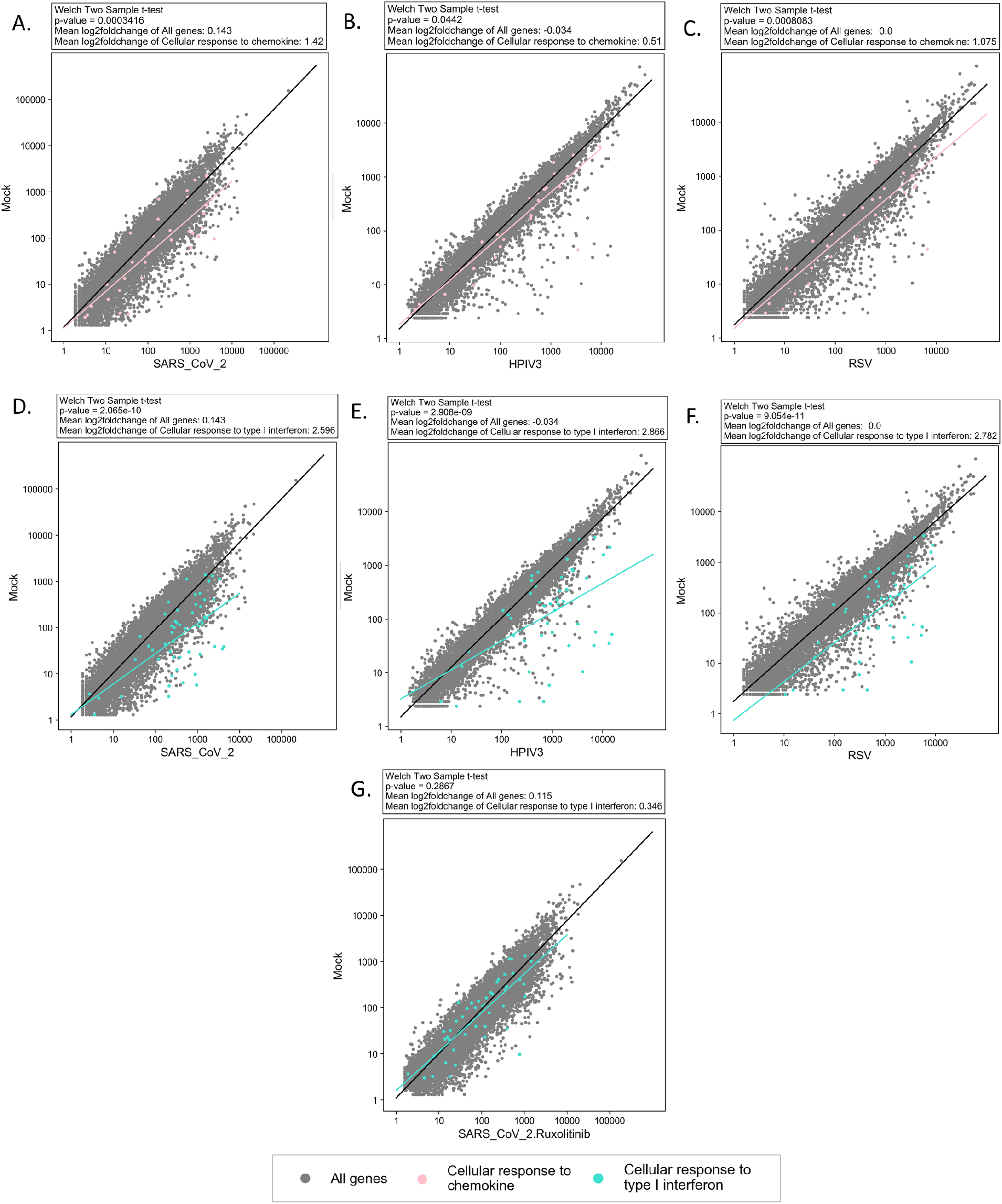
Strong chemokine and IFN-I cell response upon SARS-CoV-2, HPIV3, and RSV infection in A549 cells. A module for testing deferential expression analysis of gene groups shows that genes related to chemokine (pink) (**A-C**) and IFN-I response (turquoise) (**D-F**) are significantly upregulated, compared to all genes (grey) upon infection with SARS-CoV-2 **(A, D)**, HPIV3 **(B, E)**, and SARS-CoV-2 (**C, F**). **G**. Treatment of SARS-CoV-2 with added Ruxolitinib to block IFN-I resulted with non-significant IFN-I cell response (P-value>0.05) (turquoise), compared to all genes (grey). The black, pink, and turquoise lines are the regression lines for: all genes, chemokine related genes, and IFN-I related genes, respectively. Log scale plots present average normalized gene counts.

For a small part of the datasets of (19), for validation, we also processed the raw read data by ourselves and created the gene count tables (data is not showed). Raw RNA-seq data of human Lung adenocarcinoma cells (A549) infected with SARS-CoV-2 virus or mock, were downloaded from SRA, BioProject PRJNA615032. FastQC was used to control the read quality and trimming was performed accordingly. Reads were collapsed and first aligned to the SARS-CoV-2 reference genome version NC_045512.2 using bowtie (31). Alignment to the SARS-CoV-2 genome was made to exclude reads that are originated from the virus for further analysis, and to validate that in contrast to the mock samples, the SARS-CoV-2 samples are infected with the virus. Indeed, few thousands of reads were mapped to the SARS-CoV-2 genome, only for the SARS-CoV-2 infected samples. Unaligned reads were aligned to the human transcriptome version GRCh37 (hg19) using bowtie allowing multiple alignments and gene expression levels were evaluated by read counts. We then compared our created gene counts to the already processed counts downloaded from GEO. Although read count values were not identical, as expected due to the use of different alignment tools, the trend was the same.

## RESULTS AND DISCUSSION

### I. Description of BiSEK’s modules

To reliably analyze differential expression of genes across samples we generated BiSEK, which contains the following modules (**Figure 1A**): (1) Experimental design setup, for allocating sample information and experimental design setup (2) PaDETO, a custom algorithm for testing the data variation in relation to the desired experimental design (3) DEA based on DESeq2, for statistical parameter selection for DESeq2 run and data visualization (4) DEA of group of genes, for testing the change in expression for specific groups of genes and generating plots. The algorithms, statistical implementation and tools are lengthily described in the Materials and Methods section.

#### (1) Experimental design setup: BiSEK interactive and dynamic preprocessing workflow for multivariable data

To make BiSEK as versatile as possible, BiSEK is not dependent on any alignment tool, but uses as input a read-count table for all expressed genes or any other differential analysis needed. Samples can be uploaded to BiSEK GUI separately or by bulk (**Figure 1B**). Once uploaded, BiSEK automatically detects genes that are not being expressed in all given samples and allows the user a choice to continue with either: considering only genes that appear in all the lists, or considering all genes by filling in missing gene counts in samples with zero counts. In addition, it provides a tabular and graphical interface for the inspection of the uploaded count files (**Figure 1B-C**). This interface is updated in real time as filtering options are selected later.

In the next step the user adds any number of metadata factors that differentiate samples (**Figure 1D)**. In complex DEA study designs, where more than two conditions differing between samples are to be compared, BiSEK allows defining a multivariable data object (commonly referred to as dds) (16). The user determines the desired comparison between two values or more, within a defined factor (set by the user or created by the software) (**Figure 1E)**. For simplified analysis, the software also automatically combines the defined factors into a single factor that the user may choose to create a single condition (for example, if Genotype and Treatment conditions were defined, a Genotype_Treatment condition would be created), as recommended for certain situations by the DESeq2 vignette (16) (**Figure 1E, orange arrow**). BiSEK further supports testing interaction effects between metadata factors and automatically suggests adding interaction-terms into the design (**Figure 1E, blue arrow)**. See Methods for more information.

#### (2) PaDETO: an algorithm for testing data variation in relation to the experimental design

Scientific questions of interest, regarding comparison of gene expression-levels, inherently involve fundamental assumptions for differentiating between groups of samples. These fundamental assumptions may stem from differences between the genetic backgrounds of strains, distinct cell responses among varied conditions, and altered biological reactions that depend on the batch in which the sample was taken. Examining the expression profiles of samples that are tested for their differential expression is crucial to infer the validity of the basic assumptions on which the experimental design is founded. Specifically, it would be expected that similar profiles would be obtained for samples that are tagged under the same conditions. A common practice for examining the expression profiles of samples is a PCA. PCA, enables evaluating the correctness of an experimental design, and is an integral part of DEA outline for many researchers. However, using PCA to infer the correctness of an experimental design may be challenging and requires mathematical and statistical knowledge. Incorrect interpretation of the PCA may in some cases lead to unsuccessful data analysis and to reduced statistical power (i.e. such as a lack of detected DEG), and in more severe scenarios cause misleading results. To address this issue, we developed the novel algorithm, PaDETO (described broadly in the Supplementary Material). The PaDETO algorithm detects discrepancies between the metadata information supplied by the user about the samples, and the geometric clustering of the data obtained by PCA. PaDETO can detect data discrepancies such as an outlier sample, undefined batch effect, samples that were accidentally switched and more. The classification of discrepancies by PaDETO allows BiSEK to supply the user with actionable recommendations to improve the fit of the experimental design to the actual data behavior, leading many times to increased power the of the test for differential expression. The recommendation can then be easily applied in the BiSEK platform. **Figure 2** shows a schematic outline of the PaDETO algorithm.

To test the algorithm, we applied it to our own gene expression data from wildtype *C. elegans* worms that we know has a batch effect (30) (**Figure 3**). PaDETO was able to successfully detect the batch effect (**Figure 3A**) and the data partitioned well following batch effect variation removal (**Figure 3B**). Following batch effect detection and after defining a “Batch” condition in the condition table as recommended by the PaDETO, the power of the statistical test, to identify of differential expression, was improved. **Figure 3C, D** reflects the improvement of the power of the test as to the much larger excess of P-values in the lowest bins (P ≤ 0.10) (i.e., a larger number of rejected null hypotheses) and correspondingly, much less P-values across higher bins. Another example, using a different dataset, demonstrates the importance of normalizing the distance between samples based on all PCs variances by PaDETO, to prevent manual misleading interpretation (**Figure S2**). We further illustrate how PaDETO detects samples that were intentionally switched by us (**Figure S3**). Last, we show that PaDETO was able to detect an outlier sample (**Figure 4A**) that was not detected originally using manual PCA interpretation (18), thus leading to an improved analysis and conclusions (see below). Taking together, we concluded that PaDETO is a powerful algorithm to identify discrepancies in the data, which improves regular PCA interpretation when it is hard to consider all mathematical aspects.

#### (3) DEA based on DESeq2

The option to compare between groups containing more than 2 values within a factor, or to test interactions exists in DESeq2, but deviates from the standard DESeq2 workflow (via the DESeq2 *resultsNames* function) (16) and requires the users to construct their experimental design as a linear combination of statistical coefficients. BiSEK automatically constructs such linear combination via an easily understood GUI using the researcher’s terminology (**Figure S1A, black arrow)**. These features make BiSEK valuable for inexpert users having complicated experimental designs.

One of the greatest achievements of DESeq2 is the option to employ shrinkage of LFC values, to minimize the common problem of high LFC values that do not exhibit true expression changes between tested conditions, obtained due to low gene counts or gene counts with a high CV (8). In many studies, shrinkage can improve visualization and gene ranking. Therefore, we enable to employ shrinkage using BiSEK, with the DESeq2 *lfcShrink* function (16) (**Figure S1B)**. In case no shrinkage is desired, the main function used to create DESeq2 results, *results* (16), will be used instead (**Figure S1A)**. A detailed description of these options is described in Materials and Methods.

Visualization of the data is crucial for the validation and understanding of the results. BiSEK enables a variety of visualization options. Firstly, the DESeq2 results table is presented with an option to add normalized gene counts to all given samples for each ranked gene. Secondly, a P-value distribution plot is displayed to ensure that a large excess of P-values are in the lowest bins (P ≤ 0.10). This indicates for many rejected null hypotheses as expected in a high-powered test. In addition, plotMA, a plot showing the log fold change (LFC) against the normalized genes in average (see an example in **Figure S12**), and the result summary report are automatically displayed (16).

#### (4) DEA of groups of genes

A common practice in DEA is first to detect individual affected genes, based on thresholds on their LFC and adjusted P-value, and second to cluster the detected genes based on their function, localization, or other attributes, to identify specific pathways that are influenced. However, a pathway can be significantly affected also if the expression levels of many genes comprising it are non-significantly changed in a certain direction. These pathways will not be defined as influenced by the abovementioned scheme. To overcome this, we added a module to BiSEK to examine differential expression of groups of genes rather than individual genes. By listing one or more subgroups of genes, it is possible to visualize their expression differences versus all genes, or another sub-group of genes (see an example in **Figure 4**), and to conduct a statistical test (see Material and Methods).

### II. Upgrading previously conducted analysis, based on published data, using BiSEK

#### FUBP1 depletion causes expression enhancement of apoptotic genes and apoptotic pathways in CML cells

To test whether BiSEK can improve findings from RNA-seq data that were also tested experimentally, we concentrated on data from CML cells of Mice bone marrow (BM) from a recent paper by (18). Hoang et al. used the standard DESeq2 pipeline to test the impact of Far Upstream Element Binding Protein 1 (FUBP1) in CML Leukemia, by comparing between the RNA expression of mice that were transplanted with BM cells expressing the oncogene BCR-ABL1 and Fubp1 shRNA (Fubp1 depletion) and mice that were transplanted with BM cells expressing BCR-ABL1 and scrambled shRNA (control) (18). BiSEK algorithm, PaDETO, detected one of the Fubp1 shRNA+ samples as an outlier (Figure 4A). The PaDETO interpretation is derived from the distant location of the outlier-sample in the PCA space compared to the rest of the samples of the same type. In this case, the outlier Fubp1 shRNA+ was attributed to the cluster in which all scrambled shRNA+ samples were properly placed. It should be noted that Hoang et al. (18) performed a PCA to test the data variation in relation to the experimental design. However, the outlier was not annotated. We suggest that the outlier was not detected, because the first three PCs were plotted in 3D and the conclusions from the PCA were inferred manually. This is potentially misleading due to optical illusions caused by the projection of the 3D plot to the 2D. One more possible reason for the outlier not being detected is the deduction regarding the validity of the setup that was based on the fact that the two sample types were linearly separable in PCA space. While it is true that PCAs in which the conditions are not linearly separable are problematic, a linearly separable PCA is not sufficient for concluding that the experimental design is valid.

In accordance with the PaDETO interpretation, when removing the outlier sample and running BiSEK with the default deseq2 parameters, but defining *lfcThreshold* =1 (to identify only highly affected genes), the strength of the test was increased nearly 10 fold. Thus, the number of significantly expressed genes was increased upon removal of the outlier sample, to 344 significant genes (P-adjusted<0.05) compared to only 35 obtained if including all samples, as in Hoang et al (18) (**Tables S1-S2**).

To test whether excluding the outlier sample from the analysis affects Gene Ontology (GO) enrichment, we submitted the differentially expressed lists of genes to Gene ontology resource (32,33). When including all samples, no substantial enrichment of any GO term was obtained. However, excluding the outlier sample resulted in detection of 420 enriched biological processes with False Discovery Rate (FDR) < 0.05 (**Table S3**), indicating that PaDETO recommendation was essential in this case. Hoang et al., showed experimentally in the same system that FUBP1 deficiency enhances apoptosis (18), prompting us to search for enriched apoptotic processes in the RNA-seq data. Indeed, when excluding the outlier sample, the term “regulation of apoptotic process” (GO:0042981), for example, became significantly enriched with 37 most effected genes. As mentioned before, when including the outlier sample, no GO terms were enriched, thus obviously “regulation of apoptotic process” term was not detected as well. Therefore, the PaDETO recommendation was essential for finding highly affected processes and genes.

Next, we used an opposite approach. Knowing that FUBP1 depletion experimentally enhanced apoptosis, we used the BiSEK module that enables testing the expression tendencies of groups of genes compared to all genes, to confirm this bioinformatically. We submitted the full list of genes participating in “The regulation of apoptotic process” (GO:0042981) into BiSEK’s “DEA groups of genes” module and tested its overall expression tendencies compared to all genes. We were able to show that either inclusion or exclusion of the outliner sample, resulted in a significant expression upregulation of genes related to the apoptotic process (**Figure 4B-C**). In accordance with our previse results, showing that omitting the outlier sample leads to an increased power of the test, “The regulation of apoptotic process” became more significantly enriched if excluding the outlier (**Figure 4C**) compared to the analysis result using all samples (**Figure 4B**).

Overall, using BiSEK we were able to bioinformatically strengthen the original experimental observations. Moreover, by following PaDETO recommendation we could increase the power of the statistical test to identify highly affected genes and biological processes.

#### IFN-I response is substantial following SARS-CoV-2 infection

Currently the world is struggling against the COVID-19 pandemics, caused by SARS-CoV-2 virus. Understanding the host reaction following SARS-CoV-2 infection, which is very different from that to other familiar respiratory viruses, is crucial for the development of effective treatment and prevention of long-term sequelae. Therefore, to exploit BiSEK capabilities of grouping genes and testing the input data, we used BiSEK to reanalyze the data that were generated and investigated in the study by Melo et al (19), in which the gene expression levels following SARS-CoV-2 infection were compared to those following infections with other viruses: Influenza A virus (IAV), Respiratory Syncytial virus (RSV) and Human parainfluenza virus type 3 (HPIV3). The original study consists of series of experiments performed using different *in-vitro* and *in-vivo* models on human and ferret, from which we re-analyzed raw gene counts obtained from human models (GSE147507). We used several human sample-types classified into series, as follows: (A) NHBE cells infected with SARS-CoV-2 (series 1) or IAV (series 9) (B) A549 cells, with a vector expressing human ACE2, infected with SARS-CoV-2 high-multiplicity of infection (MOI) (series 16) and low-MOI (series 6) or RSV and HPIV3 (series 8) (C) post-mortem Lung samples of COVID-19 patient (series 15). Using BiSEK, we analyzed each series separately as was originally done (19).

Cytokines are a large group of polypeptides or glycoproteins, produced mainly by immune cells, that responsible for cell signaling to evoke an immune response. Cytokines include: chemokines, interferons (IFNs), interleukins (IL), lymphokines and tumor necrosis factors (TNFs) (34). It was reported by Blanco-Melo et al. (19) that SARS-CoV-2 evokes a strong chemokine inflammatory response, which varies from the response upon other virus infections. It was also noted that generally, IFN-I and -III expression levels are low upon SARS-CoV-2 infection and that a mild IFN-stimulated genes (ISG) response is obtained. However, a statistical test inferring the differences in expression levels of group of genes, for each virus treatment versus mock, was not conducted. We therefore used BiSEK’s “DEA of group of genes” module, to test the expression tendency of genes related to chemokine and ISGs activity among all series of samples mentioned above. We selected groups of genes to reflect chemokine, IFN-I, and IFN-II cellular responses, and tested their expression level changes following each virus treatment. These genes belong to the following GO categories: “cellular response to chemokine” (GO:1990869), “cellular response to type I interferon” (GO:0071357), and “cellular response to interferon-gamma” (GO:0071346). We also tested the general response of cytokine related genes (GO:0034097), which includes among others, all the genes from the above GO terms.

We note that since the raw read counts for many IFNs was zero, for both mock and treated samples, we could not evaluate their differential expression levels between samples, relying solely on the RNA-seq data. These genes are expected to be expressed under standard conditions (35). Therefore, the lack of IFNs expression in mock samples, may be a result of alignment difficulty. For this reason, we evaluated the interferon response via ISGs that are transcriptionally activated by interferon genes (35).

We first run the BiSEK’s module “PaDETO”, to test the fitness of NHBE infected-cells data. It was indicated by PaDETO that the data fit well to the experimental design for SARS-CoV-2 and their corresponding mock treated sample (**Figure S4A**). Also, there is a clear separation between IAV samples and their corresponding mock samples, despite a notable unexplained variation between IAV samples that were split into two clusters (**Figure S4B**). Next, we run DEseq2 via BiSEK interface with the default parameters to test the expression levels upon each virus infection (**Tables S4-S5)**. IAV infection resulted in a much larger number of significantly DEG compared to SARS-CoV-2 (2452 versus 412; P-adjusted value<0.05). As indicated in the original study (19), a large part of the DEG was not common (only 105 common genes). This may indicate a real difference between the two infections, but since each test was performed separately, this may also be the result of other confounding variables. Despite this difference, both viruses caused an extensive cytokine response compared to their corresponding mock controls (**Figure S5A-B;** P-values**=** 1.07E-13 and 6.44E-11 for SARS-CoV-2 and IAV, respectively), indicating for a massive general inflammatory reaction.

Following the indication for a general inflammatory response in NHBE, we wished to interrogate the difference in response of specific immunomodulating agents (chemokine, IFN-II, and IFN-I) that are included in the cytokine group. We therefore tested the differential expression tendency of genes belong to (1) chemokine, (2) IFN-II, and (3) IFN-I cellular reaction, using BiSEK “DEA of group of genes” module (**Figure 5**). Our results indicate that: (1) SARS-CoV-2 and IAV infections resulted in a significant chemokine response (P-value = 1.35E-03, and 3.56E-03 respectively), consistent with the original analysis (19). (2) The cellular response to IFN-II was substantial as well, supporting Blanco-Melo et al.’s findings (19) (**Figure S6A-B;** P-values**=** 1.97E-03 and 9.33E-03 for SARS-CoV-2 and IAV, respectively). (3) In contrast to Blanco-Melo et al. reporting that IFN-I response is diminished in NHBE cells upon SARS-CoV-2 infection (19), our analysis shows that IFN-I response in NHBE is similar to that of the chemokine (mean log2FoldChange values 0.635 and 0.593, respectively) and even more significant (P-values = 4.74E-07 for IFN-I compared to 1.35E-03 for chemokine). The IFN-I cell responses to SARS-CoV-2 and IAV were similar in respect to ISGs (P-value =1.12E-06 for IAV and P-values = 4.74E-07 for IFN-I), but the reaction to IAV was stronger based on mean Log2FoldChange values ((0.635 - (−0.015))= 0.65 for IFN-I compared to (1.407 - 0.042)= 1.365 for chemokine). The statistical evaluation of the IFN-I reaction as a group of genes is probably the cause of why our findings are not in full agreement with the conclusions by Blanco-Melo et al (19).

To test whether the observations obtained for NHBE cells can be reproduced using other sample types, we next tested in a similar manner the immune response in A549 cells (with a vector expressing human ACE2) following infection by several virus types. We first used PaDETO to test the data behavior for A549 cells.

There were no inconsistencies in the data for SARS-CoV-2 and the corresponding mock samples (series 16) (**Figure S4C**). However, when testing RSV and HPIV3 treatments compared to their corresponding mock samples, one of the mock samples was marked as an outlier **(Figure S4D)**. Indeed, the power of the test was increased when removing the outlier sample (**Figure S7**). We therefore removed the outlier sample when performing all analysis steps listed below.

We used the default DESeq2 parameters and tested the high-MOI treatment of: RSV, HIPV3, SARS-CoV-2, and SARS-CoV-2 with added Ruxolitinib (to block IFN-I response), each versus its corresponding mock sample, as originally done (19). We then utilized the BiSEK “DEA of group of genes” module as described above to test the tendency of genes that represent a general cytokine response, in each of the treatment types, for A549 cells. In all cases, a robust cytokine response was achieved, indicating a general strong immune activation (**Figure S5**; P-values= 4.04E-13, 2.20E-16, and 2.20E-16 for SARS-CoV-2 and RSV and HIPV3, respectively), as was also shown for NHBE cells.

Concordant with the same strategy described above for NHBE cells, we next tested the differential expression tendency of genes belonging to (1) chemokine, (2) IFN-II, and (3) IFN-I cellular reaction, to highlight the difference in response to specific immunomodulating agents under the cytokine group, upon exposure to different viruses.

(1) The cell response to chemokines was significant in all cases, while for SARS-CoV-2 and RSV a slightly stronger result was presented compared to HIPV3 (**Figure 6A-C**; P-values= 3.40E-04, 8.00E-04, and 4.42E-02 respectively). Similar to the observations for NHBE cells, a strong IFN-I response was obtained for all virus kinds, including SARS-CoV-2 infection in high-MOI (**Figure 6D-F**, P-values= 2.07E -10, 2.91E-9, and 9.05E -11 for SARS-CoV-2, HPIV, and RSV, respectively). (2) IFN-II reaction was obtained as well against all virus types (**Figure S6C-E;** P-values= 1.90E-07 and 9.39E-09, and 1.15E-10 for SARS-CoV-2 and HIPV3, and RSV respectively). (3) Strikingly, when testing the IFN-I response upon SARS-CoV-2 infection, but with the addition of Ruxolitinib to block IFN-I response (series 16), the tendency of IFN-I related genes was similar to all genes (**Figure 6G**, non-significant P-value). Weaker IFN-I response upon SARS-CoV-2 infection following Ruxolitinib addition was also observed by Blanco-Melo et.al (19), supporting our obtained trend (Figure S1H in (19)). The lack of a considerable IFN-I response in the presence of Ruxolitinib serves as a negative control and corroborates our findings supporting the strong IFN-I response after SARS-CoV-2 infection.

It was claimed in the original study that the cell immune response, originated particularly from IFN-I activation, is prevented by an antagonist under low-MOI SARS-CoV-2 infection, but not under high-MOI, for which the antagonism is inefficient (19). We, therefore, tested the immune response upon low-MOI SARS-CoV-2 infection (series 6). Our analysis illustrates that indeed a much weaker IFN-I response is obtained following low-MOI SARS-CoV-2 infection, compared to high-MOI (**Figure S8A;** P-value>0.05). However, in the case of low-MOI a general cytokine and IFN-II cell response are lacking as well (**Figure S8B-C;** P-values>0.05), suggesting that overall, the immune system reacted weakly upon low virus titration. The group of chemokine is the only group among the others that reach a statistically significant response (**Figure S8D;** P-value=1.02E-02). Although statistically significant, the chemokine response obtained for low-MOI was weaker than that for high-MOI, for both A549 (P-values = 3.40E-04) and NHBE cells (P-value = 1.35E-03) (**Figure 5A**; **Figure 6A**). We therefore suggest that the relatively weak IFN-I response in low-MOI treated cells is concordant with a general weak immune response, thus not reflecting imbalanced host response in this case. Additional studies of low-MOI infections should be performed to achieve further conclusions.

To infer our findings for clinical samples, we tested the immune system activity in lung tissues originated from post-mortem COVID-19 patients. Similarly to the results of our analysis of the *in-vitro* samples, a substantial general immune response was accrued in COVID-19 patients, indicated by a significant cytokine response (P-value=3.86E-04). In addition, all specific immunomodulating subgroup agents (chemokine, IFN-I, and IFN-III) significantly upregulated, while IFN-I and IFN-II related genes achieved a stronger response compared to chemokine (**Figure S9;** P-values= 5.41E -6, 6.73E -9, and 7.09E -3 for IFN-I, IFN-II, and chemokine, respectively).

Overall, up to this point, our results suggest that SARS-CoV-2 infection in humans leads to an extensive immune response, that includes a robust IFN-I activity. The difference in conclusions between Blanco-Melo’s (19) and ours, may be the result of different statistical analysis strategies as well as the use of distant methods to evaluate DEG group response, foremost among them is the BiSEK module to test the expression tendency of sub gene groups. Also, by removing an outlier sample from one of the datasets, we might increase the variance between the two analyses. Within this context, it is encouraging to note that our results are supported by other recent studies in the field examining IFN-I response upon different levels of disease severity and varied sampling time points. V’kovski et al., shows a substation IFN-I and ISGs expression upon SARS-CoV-2 infection (36), Lee et al., shows how IFN-I response is enhanced specifically under severe COVID-19 and suggests that IFN-I reaction plays a key role in provoking the typical COVID-19 severe inflammation response (37), and Zhou et al., reports enhanced expression of ISGs, regardless of the severity of COViD-19 disease (38). It should be noted that one more study reported impaired IFN-I response (39). However, this study (39) only partly supports the findings of Blanco-Melo et al. (19), since its conclusion of low IFN-I response referred to severe COVID-19 cases, while in Blanco-Melo et al. impaired IFN-I response was shown for *in vitro* low-MOI treatment of SARS-CoV-2, but the opposite was true for COVID-19 patients. Finally, it was explicitly mentioned in a recent review that there are contradictory reports regarding the intensity of IFN-I reaction in COVID-19 patients (40). There is a prevalent speculation that the origin of the difference in the extent of IFN-I response is the disease severity (37,40). Based on our interpretation of Blanco-Melo et al.’s data and the indication from others that IFN-I response is substantial under different severity levels of the disease, we suggest that some of the differences are due to the usage of distinct analysis and statistical methods.

It was further found in the original study that a specific chemokine signature, relying on CXCLs, IL-6 and IL-1, distinguish the host SARS-CoV-2 response from that of other viruses (18). While we do not deny a signature of that type, based on our analysis it could not be tracked (**Figure S10**). The fact that not all CXCLs, IL-6 and IL-1 signature-related genes were expressed in a sufficient extent to be analyzed for all given samples, makes drawing conclusions difficult.

As previously mentioned, we analyzed using BiSEK each GEO series separately as was originally done (19). Within this context, it should be mentioned that the experimental differences arising from setting each experiment separately, cannot be addressed in the statistical design (taking into consideration all variation factors would result in a non-full rank model matrix, see (16)). Therefore, directly comparing the host responses for a given sample type following each virus infection, using the same corresponding mock as a control for all experiments, would be beneficial.

#### TLR3, related to IFN-III production regulation, may be responsible for the low IFN-III response upon SARS-CoV-2 infection

To further strengthen our results, we performed a GO analysis, using the high-MOI A549 samples, on the most highly affected genes. For this purpose, we used a stricter approach for selecting differentially expressed genes. Namely, rather than running the DESeq2 default, we used the same data as before (serious 16 and 8) but defining lfcThreshold =1 option using BiSEK (The exported DESeq2 analysis results are presented in **Tables S6-S8**). As described above, this strict test would enable highlighting the most affected genes, by incorporating the desired null hypothesis into the DESeq2 model. To test differences in GO enrichment of significant genes, we used the Gene Ontology resource (32,33). For more accurate analysis we used all expressed genes in each dataset as the reference list.

Not surprisingly, the GO analysis shows a substantial enrichment for processes related to the immune system, including processes related to cytokine response, immune defense response, virus infection response and more (**Tables S9-S11**). Processes related to IFN-I regulation were enriched following exposure to all virus types, including SARS-CoV-2, corroborating our previous finding that SARS-CoV-2 triggers a strong IFN-I response. Processes related to interferon-gamma and interferon-betta were also enriched for all infection types. But a more curios result was obtained for type III interferon response, which was not tested in previous sections. We found that a process related to regulation of type III interferon (IFN-III) production (GO:0034344), which includes only three genes, was enriched upon RSV and HIPV3 infection, but not upon SARS-CoV-2 infection. (Tables S9-S11). This observation is in line with the general findings of the original study (18), that IFN-III response is weaker upon SARS-CoV-2 infection compared to other respiratory viruses.

To further interrogate the obtained GO result, we used the BiSEK module: “DEA of groups of genes”, for testing the average tendency of the 3 genes in the category of “regulation of type III interferon (IFN-III) production” (GO:0034344), compared to that of all genes. Indeed, a significant different tendency was obtained for RSV and HPIV3 treatments but not for SARS-CoV-2 (P-values: 2.55E-02, 1.01E-02 and >0.05, respectively, **Figure 7**). Although the test related to SARS-CoV-2 did not reach statistical significance, the apparent difference between the mean log2FoldChange values, for all genes versus IFN-III related genes (0.143 and 3.632, respectively), implies that regulation of IFN-III related-genes are triggered (**Figure 7C**).

**Figure 7:**
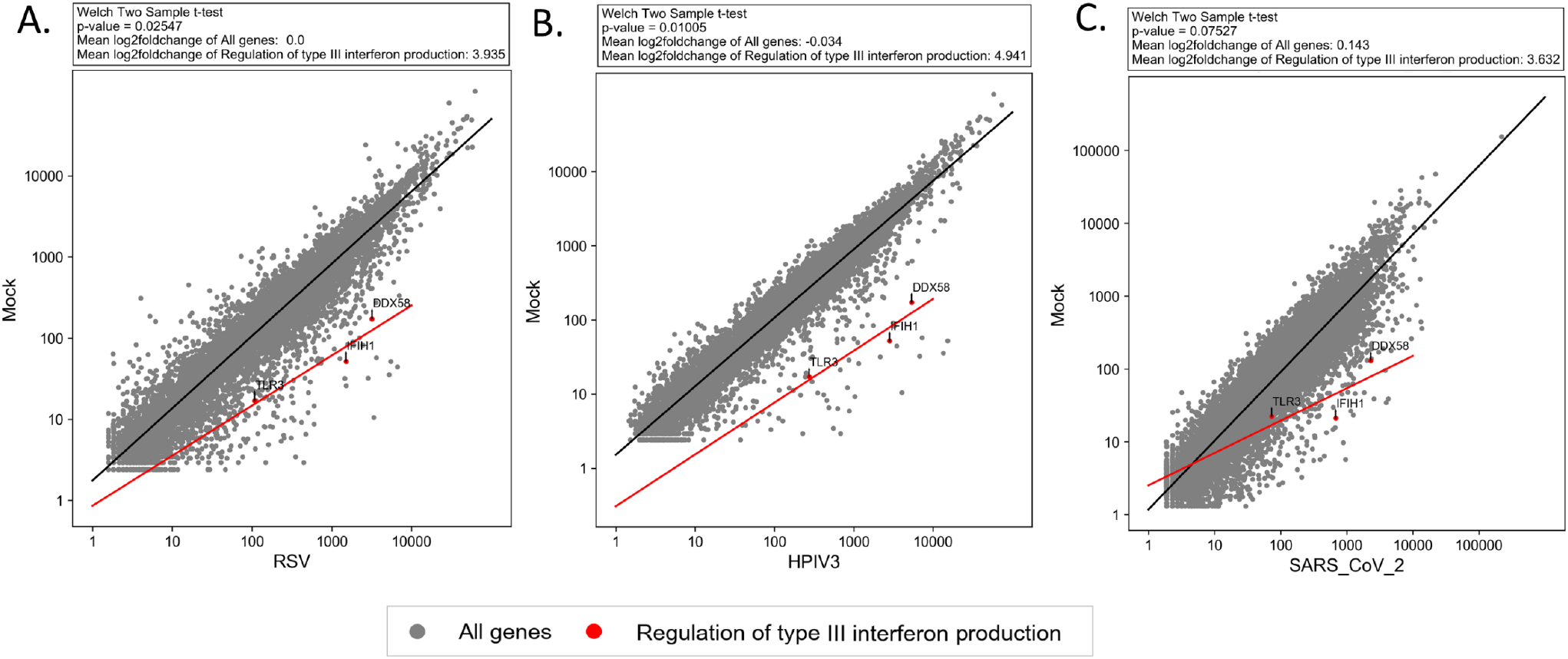
Genes involved in regulation of type III interferon production significantly differentially-expressed upon RSV and HIPV3 infections but not upon SARS-CoV-2, in A549 cells. The BiSEK module for testing deferential expression analysis of gene groups shows that genes involved in the regulation of type III interferon production (red) are significantly upregulated, compared to all genes (grey) upon infection with RSV **(A)** and HPIV3 **(B)**. The test does not reach statistical significance for SARS-CoV-2 infection **(C)**. The Log scale plots presenting average normalized gene counts. The black and red lines are the regression lines for all genes and regulatory IFN-III genes, respectively.

The above result intrigued us to go over each of the three genes participating in the “regulation of type III interferon production” term (GO:0034344): TLR3, DDX58 and IFIH1, and to examine their expression level changes (expressed by the p-adjusted value of each individual gene as defined according to DESeq2 analysis) upon the infections with different viruses. Interestingly, based on the strict test all three genes were significantly differentially expressed upon RSV and HIPV3 infections, however, upon SARS-CoV-2 infection, while DDX58 and IFIH1 were highly significant, TLR3 was not (**Table 1**; **Table S6-S8**).

**Table 1:** P-adjusted values according to DESeq2 analysis for genes participating in of regulation of type III interferon production (GO:0034344)

|  | P-adjusted values |  |  |
| --- | --- | --- | --- |
|  | RSV | HIPV3 | SARS-CoV-2 |
| ***DDX58*** | 1.09E-68 | 8.74E-106 | 4.09E-151 |
| ***IFIH1*** | 2.25E-65 | 9.77E-101 | 8.26E-76 |
| ***TLR3*** | 6.74E-05 | 5.70E-16 | >>0.05 |

Next, we tested the expression tendency of the same group of genes in another cell type, NHBE, upon SARS-CoV-2 and IAV infections and in COVID-19 clinical samples. In agreement with our findings for A549 cells, the group did not reach statistical significance in samples of any type infected with SARS-CoV-2, in contrast to upon IAV infection (**Figure S11**). In particular, the change in expression of TLR3 between treatment and mock seems to be lower after SARS-CoV-2 infection compared to other virus types, when comparing against samples within the particular cell type (Log2FoldChange values in NHBE cells: -0.391 for SARS-Cov-2 and 1.131 for IAV; Log2FoldChange values in A549 cells: 1.660, 2.721, and 4.07 for SARS-CoV-2, RSV, and HIPV3 respectively) (**Tables S4-S8**). Collectively, our results suggest that a specific gene related to IFN-III production regulation, rather than a complete process is less upregulated upon SARS-CoV-2 infection compared to other viruses. This suggestion should be further tested. Overall, our analysis supports the findings of (19), regarding the relatively weak, although not completely lacking, IFN-III response upon SARS-CoV-2 infection.

All three genes participating in regulation of type III interferon production have an important role in innate immunity and help to detect, directly or indirectly, viral nucleic acids (41). The most interesting factor out of that group, TLR3, is a member of the Toll-like receptor family, which is one of the central protein classes to recognize infection and trigger the innate immune response (42). Particularly, TLR3, as well as TLR7, 8, and 9, are found in the endosomes and have an important role to detect nucleic acids originated from viral infections (43,44). In mammalian, TLR3 is known to recognize a viral double-stranded RNA (sdRNAs) that is produced following an intrusion of positive-strand RNA viruses, but less upon negative-strand RNA viruses, and to trigger the activation of NF-kappaB (NF-κB) and IFN-I. (45,46). Among the viruses tested in this study, SARS-CoV-2 is the only positive-strand RNA virus, while RSV, HIPV, and IAV are negative-strand RNA viruses. Therefore, it is very surprising, that there were no major changes in the expression of TLR3 specifically after SARS-CoV-2 infection, where TLR3 activation is expected to be prominent (47).

There was an indication for an increased TLR7 and TLR8 activation in severe COVID-19 patients, leading to a high Bruton tyrosine kinase activity (48). However, to our knowledge, no prior observations can explain our findings regarding weak induction of TLR3 upon COVID-19 infection. Further experimental work is still required to validate our findings and to understand the possible mechanisms by which a weak IFN-III related response is observed.

To summarize, our analysis suggests that SARS-CoV-2 infection evokes a substantial immune response, which is balanced considering IFN-I, IFN-II, and chemokine responses. Under the limitation of the analyzed data, our results could imply that the IFN-III response evoked by SARS-CoV-2 is weaker compared to that triggered by other virus types, possibly due to lower expression of TLR3. These results motivate further studies into the regulation of IFN-III type host response under SARS-CoV-2 infection.

## DATA AVAILABILITY

BiSEK is available as a web-interface. It is also provided as a software package that can be easily installed locally.

## ACKNOWLEDGEMENT

We thank Benjamin TenOever and Daniel Blanco-Melo for providing their datasets before publication and for sharing information about the methods. We gratefully acknowledge Guy Horev, a member of the Technion Bioinformatics Knowledge Unit, for helpful discussions. We thank Berta Eliad, Clara Frydman and Alla Fishman for comments on the manuscript.

## AUTHOR CONTRIBUTION

R.H, D.L, Y.F and A.T.L. conceived and designed the study. R.H, D.L, and Y.F implemented and developed BiSEK. Y.F carried out the interface design, with contributions from N.F. R.H analyzed the data. A.T.L supervised the work. R.H, D.L and A.T.L wrote the manuscript with input from all authors.

## SUPPLEMENTARY MATERIAL

### Supplementary Figures

**Figure S1:**
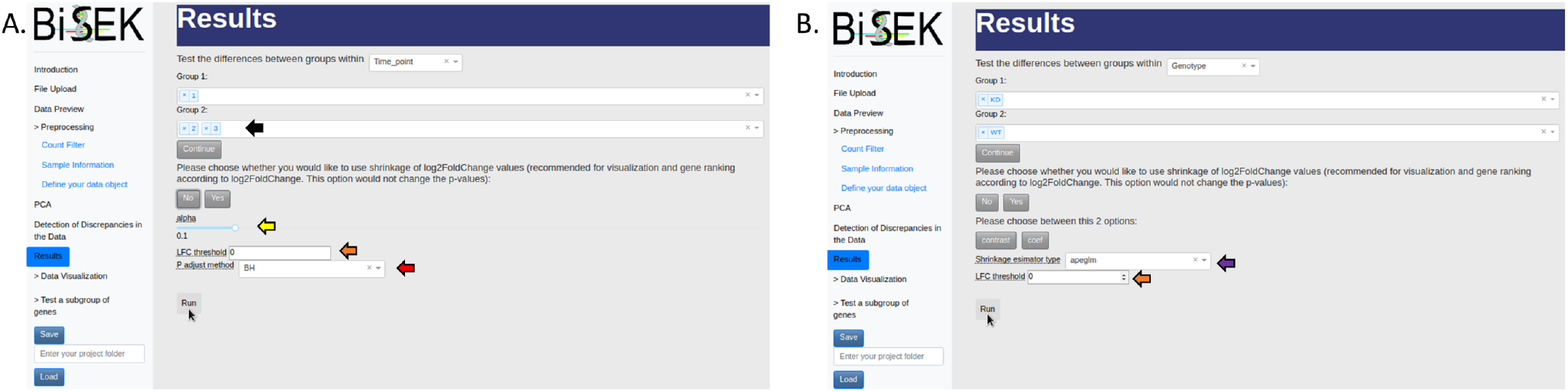
Flexible and customizable setup for running DESeq2. **(A)** No shrinkage. Black arrow, illustrates that more than one value within a condition can be selected for the statistical test. Yellow arrow, permits defining a significance level. Orange arrow, controlling lfcThreshold (if greater than 0, the P-values will be calculated for log fold change values that are greater in absolute value than the threshold). Red arrow, defining the preferable method for adjusting p-value **(B)** Shrinkage is being employed. Purple arrow, selecting shrinkage estimator type. Orange arrow, controlling lfcThreshold.

**Figure S2.**
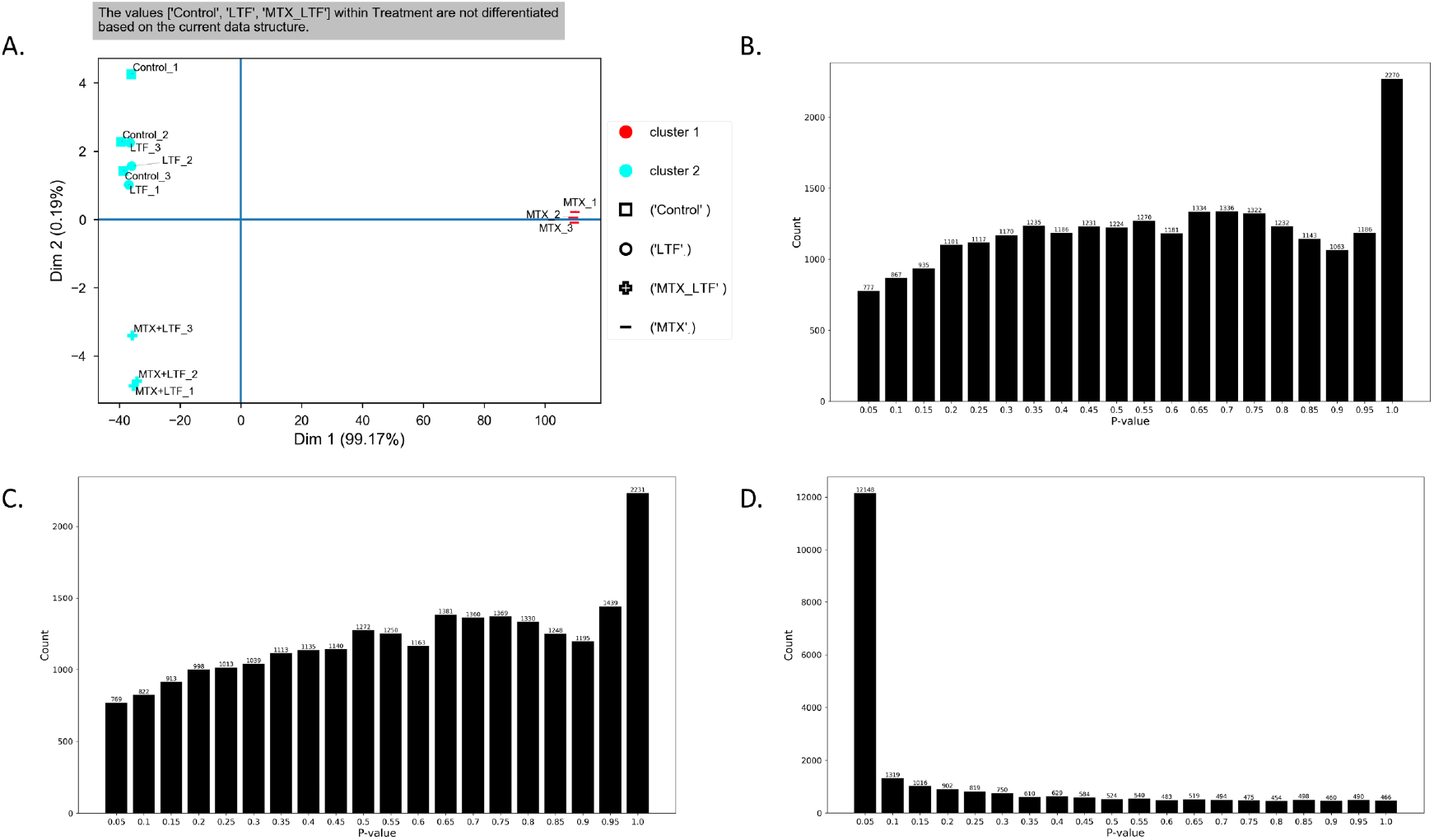
PaDETO interpretation points that some groups that were defined under the same condition are not differentiate based on a given data structure. Data was taken from GEO, accession number GSE140066. Caco-2 cells were treated with methotrexate, (MTX) lactoferrin (LTF), lactoferrin with methotrexate (MTX+LTF), and carrier fluid (control) in 3 replicates each. Colors represent the clusters by which PaDETO partitioned the samples based on their variation. Shapes represent the expected partition as defined by the user. At first glance, it seems that the data can be divided into 3 main treatment groups based on the PCA: 1) LTF and control (not differentiate from each other) 2) MTX+LTF 3)MTX. But, PaDETO’s PCA interpretation indicates that LTF, LTF+MTX and control groups (groups 1+2 described above), within treatment condition, are not separated from each other based on the currant data structure (although not all with the same shape, all colored in light blue). The PaDETO interpretation is due to the low variance of the samples in the y axis, that taken into consideration when normalizing the distance between samples based on all PCs variances. Indeed, the P-value distribution obtained by comparing the expression levels of the control to LTF+MTX (**B**) or LTF to LTF+MTX (**C**) indicates that at most, a small fraction of hypotheses are truly non-null. Therefore, there are no genes that are differentially expressed between the compared group. For comparison, when testing MTX group, versus the: control, LTF and LTF+MTX, the strength of the test is very high, i.e results with many rejected null hypothesis (**D**). This matches PaDETO’s observation that MTX is well separated from the other groups. **B-D** present p-value histograms.

**Figure S3:**
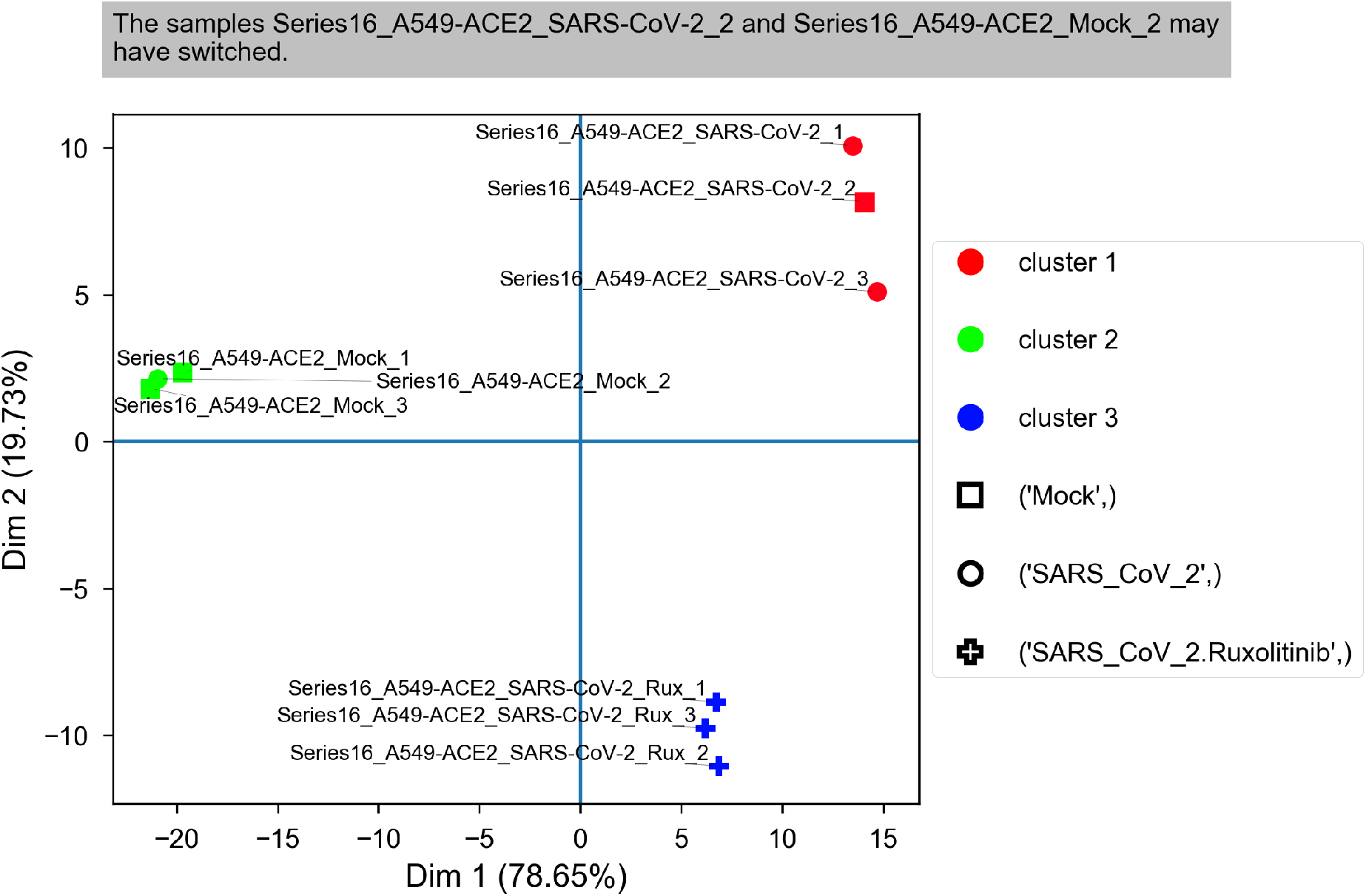
PaDETO identifies switched samples. Two samples were intentionally switched to test PaDETO ability to identify switched samples, based on data from GEO GSE147507, series 16. Colors represent the clusters by which PaDETO partitioned the samples based on their variation. Shapes represent the expected partition as defined by the user. PaDETO correctly identified that one sample from the Mock series and one sample from the SARS_CoV_2 series were switched.

**Figure S4:**
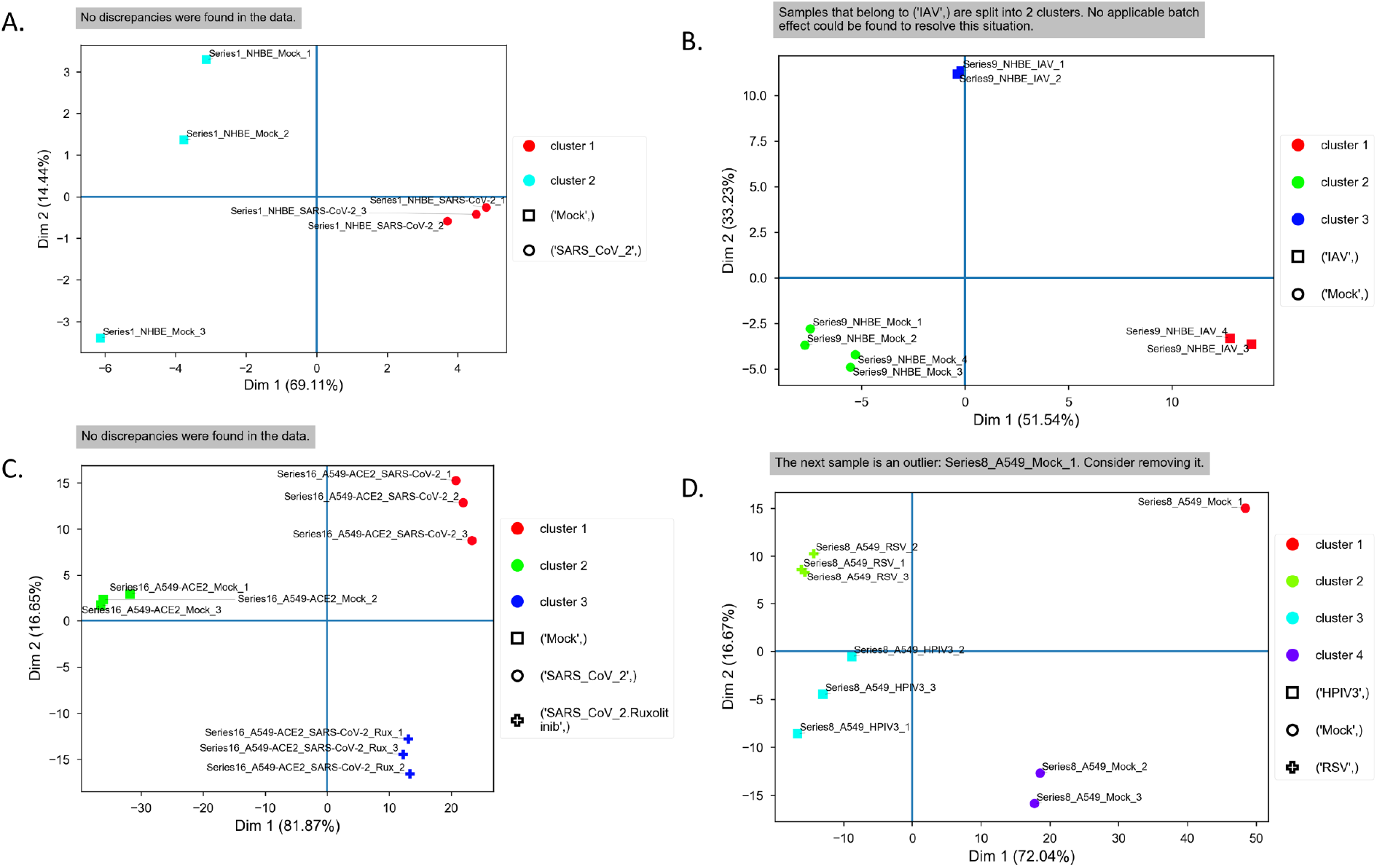
PaDETO’s PCA interpretation of datasets related to GEO GSE147507. Colors represent the clusters by which PaDETO partitioned the samples based on their variation. Shapes represent the expected partition as defined by (1). (**A)** PaDETO declares no discrepancies in the data for NABH cells that were treated with SARS-CoV-2 or Mock. (**B)** IAV samples are split off into two clusters. It is indicated by PaDETO that this issue cannot be addressed in the statistic design. Despite that, a clear separation between mock and IAV samples was obtained leading to a successful test. **(C)** No discrepancies were found in the data for A549 cells that were treated with SARS-CoV-2, SARS-CoV-2 + Ruxolitinib, and Mock. (**D)** PaDETO’s PCA interpretation pointed out one of the mock samples as an outlier, for A549 cells that were treated with RSV, HPIV3, and Mock.

**Figure S5:**
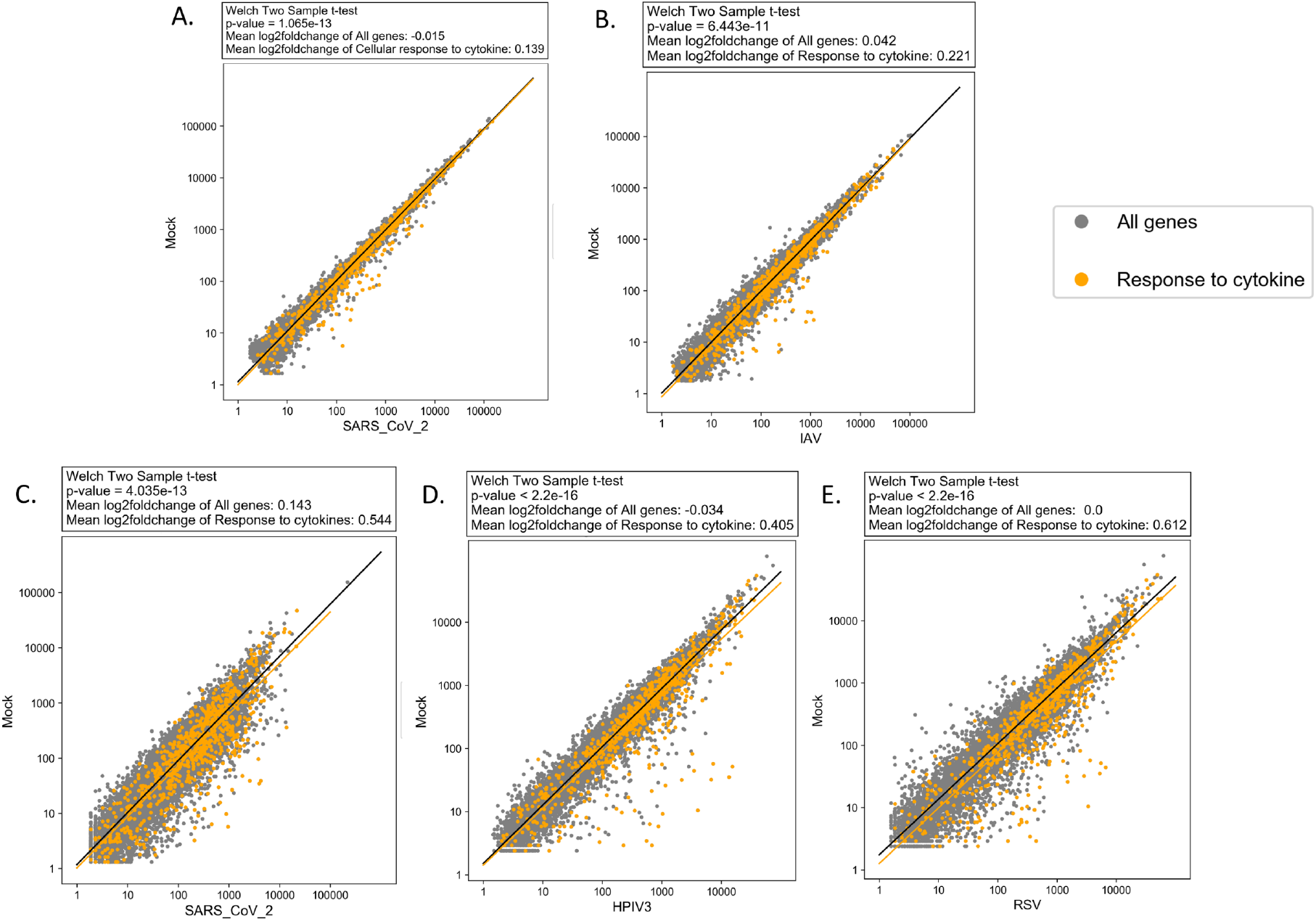
Strong cytokine response following different virus infections. Cytokine related genes (orange) are significantly upregulated, compared to all genes (grey) upon SARS-CoV-2 (**A**) and IAV (**B**) infection in NHBE cells and upon SARS-CoV-2 (**C**), HPIV3 (**D**), and RSV (**E**) in A549 cells. The black and orange lines are the regression lines for all genes and cytokine related genes, respectively. Log scale plots present average normalized gene counts.

**Figure S6:**
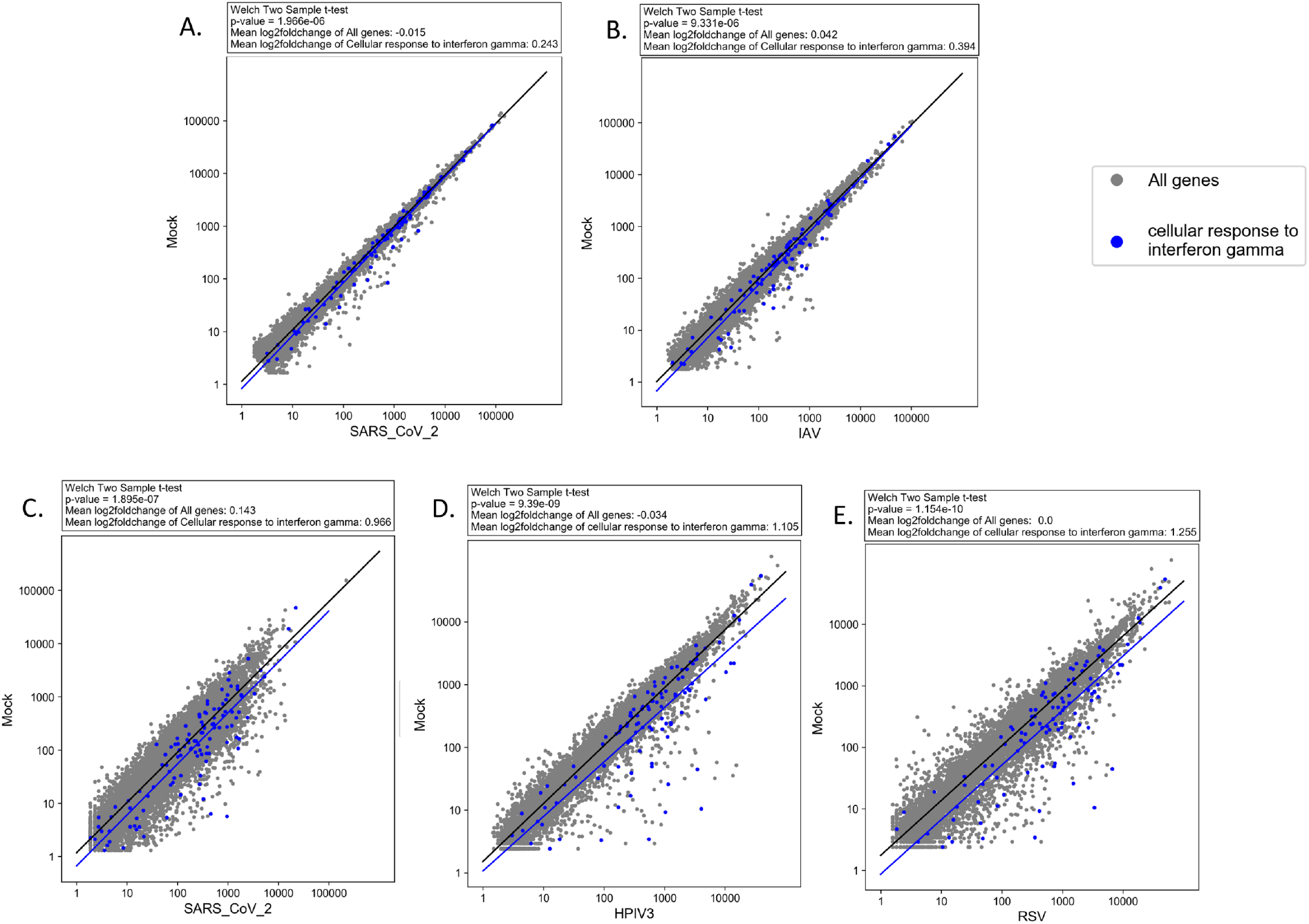
Strong IFN-II cell response following different virus infections. IFN-II related genes (blue) are significantly upregulated, compared to all genes (grey) upon SARS-CoV-2 (**A**) and IAV (**B**) infection in NHBE cells and upon SARS-CoV-2 (**C**), HPIB3 (**D**), and RSV (**E**) in A549 cells. The black and blue lines are the regression lines for all genes and IFN-II related genes, respectively. Log scale plots present average normalized gene counts.

**Figure S7:**
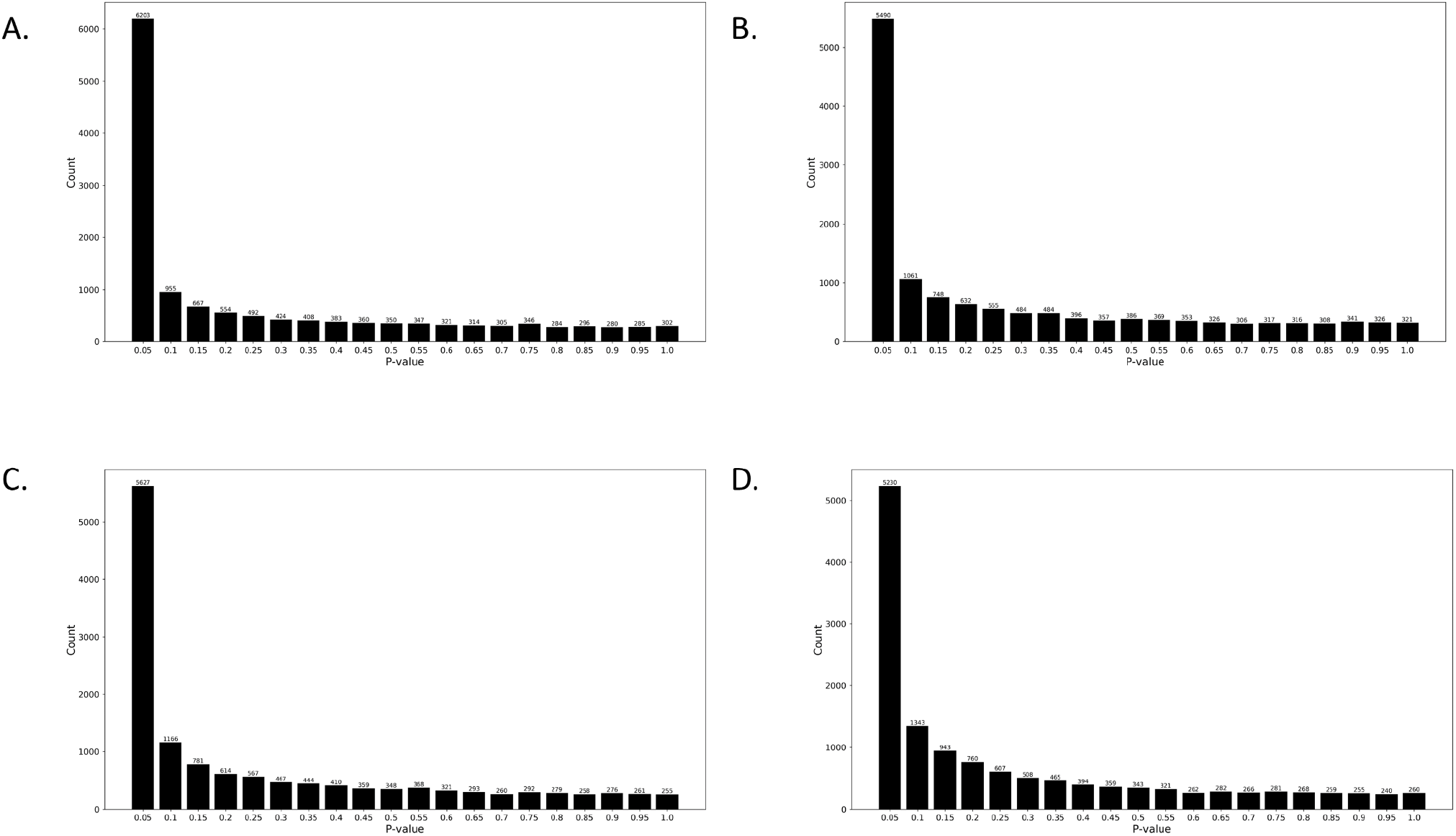
Omitting mock-1 outlier sample, according to BiSEK recommendation, resulted in increased power of the statistical test. P-value distribution plots show a larger excess of P-values in the lowest bins (P ≤ 0.10) when the test that did not include the outlier sample (A-B) compared to all samples (C-D), thus indicates the presence of a larger number of rejected null hypothesis without the outlier sample. RSV versus Mock test results are shown in A,C. HIPV3 versus Mock test results are shown in B,D.

**Figure S8:**
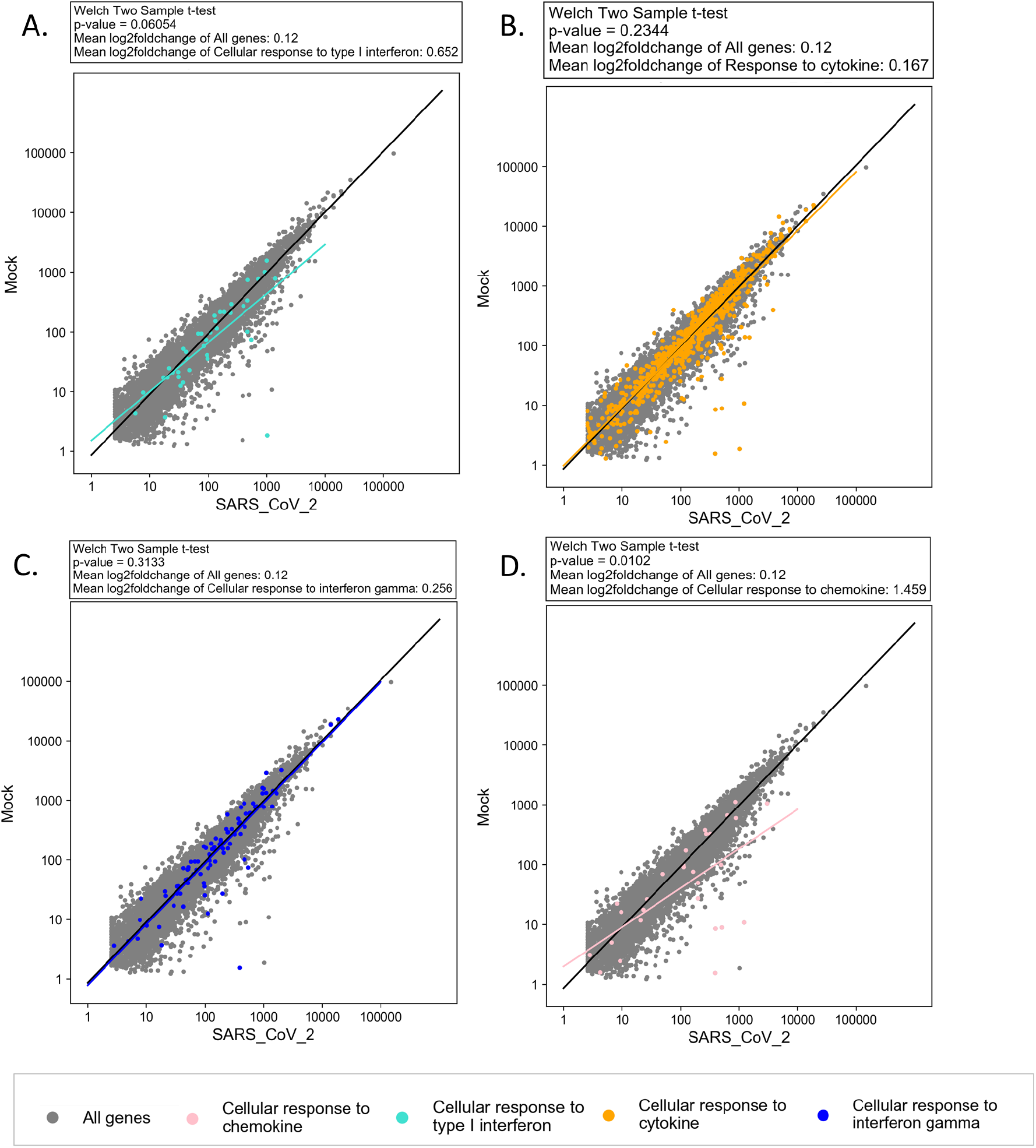
Weak immune response is accrued following infection with low-MOI SARS-CoV-2. Non-significant IFN-I (**A**), cytokine (**B**), and IFN-III (**C**) cell response following low-MOI SARS-CoV-2 infection in A549 cells. (A-**C**) IFN-I (turquoise), cytokine (orange), and IFN-II (blue) group tendency is compared to all genes (grey). **(D)** The upregulation tendency of chemokine related genes reaches statistical significance. Chemokine (pink) group tendency is compared to all genes (grey). The black and colorful lines are the regression lines for all genes and each tested sub-group, in respective to the dots color. Log scale plots present average normalized gene counts.

**Figure S9:**
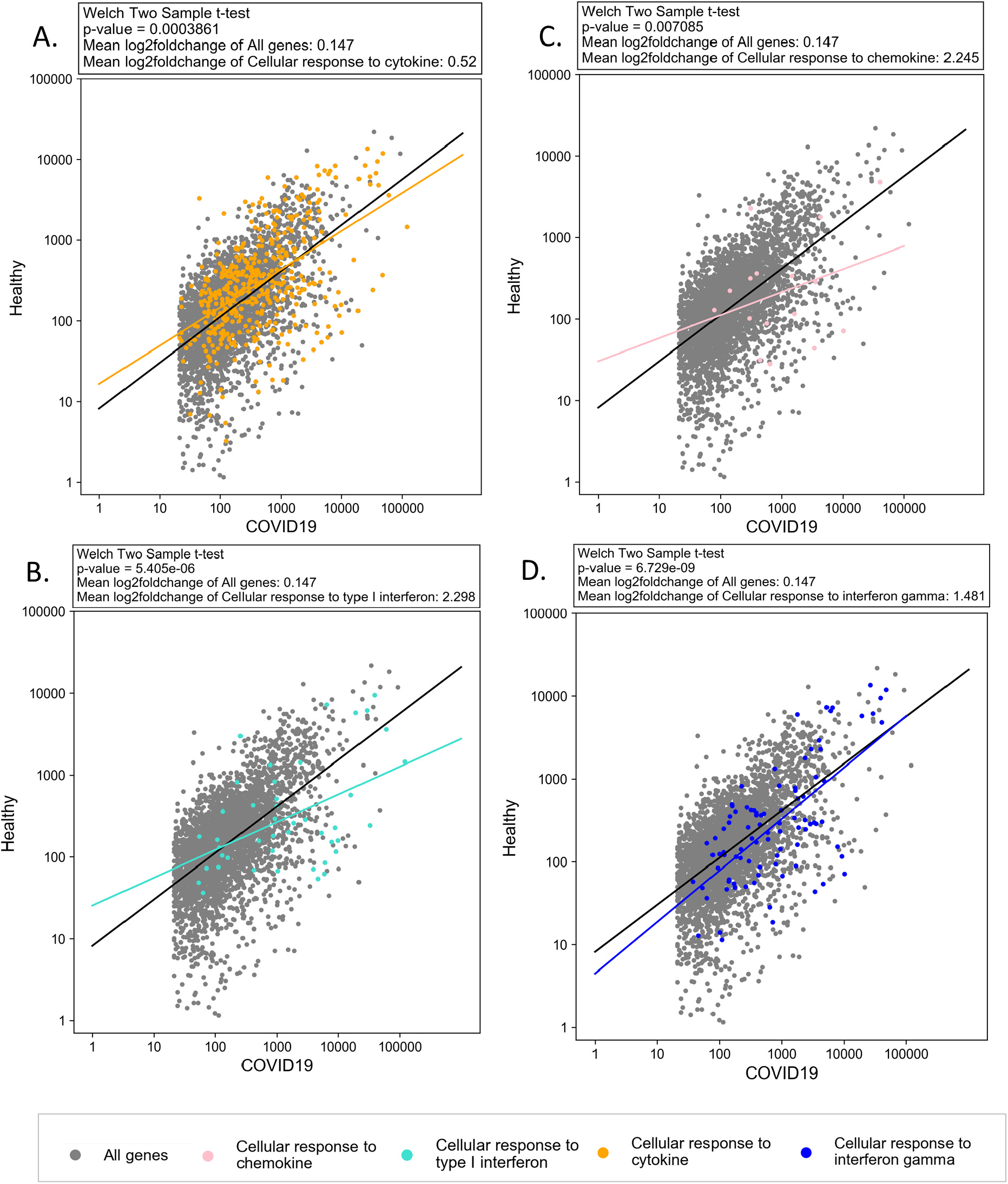
Substantial immune response in clinical COVID-19 Patients. cytokine **(A)**, IFN-I **(B)**, chemokine **(C)** and IFN-III **(D)** related gene groups are upregulated in COVID-19 patients compared to healthy tissue samples. The group tendency of cytokine (orange), IFN-I (turquoise), chemokine (pink) and IFN-II (blue) is compared to all genes (grey). The black and colorful lines are the regression lines for all genes and each tested sub-group, in respective to the dots color. The missing counts for low counts are due to data filtering (see Methods)

**Figure S10:**
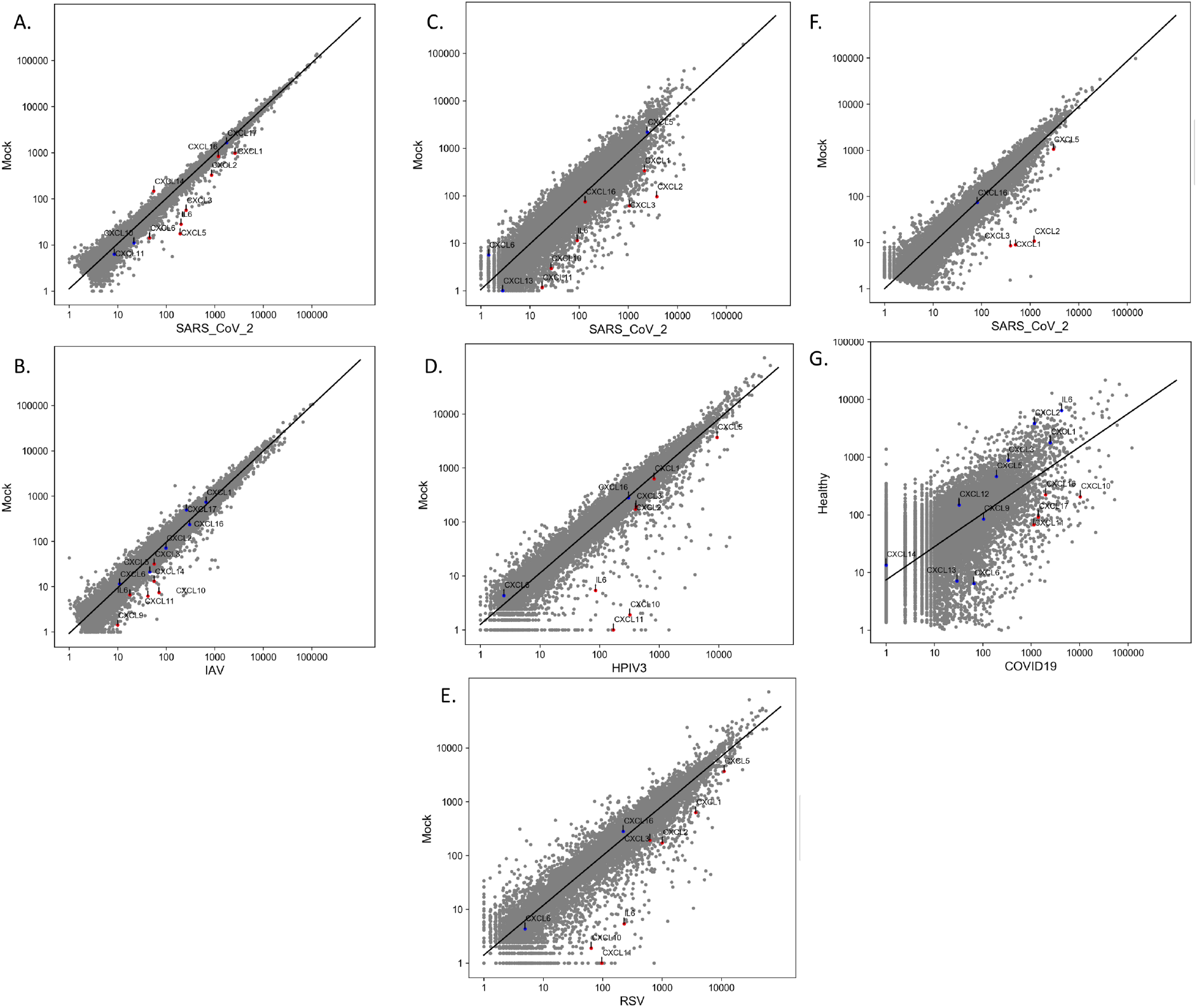
CXCLs, IL-6 and IL-1 expression signature upon different infection types. CXCLs, IL-6 and IL-1 are labeled on log scale plots that present average normalized gene counts upon SARS-CoV-2 **(A)** and IAV (**B**) infections in NHBE cells; upon SARS-CoV-2 high-MOI **(C)**, HPIV3 **(D)**, and RSV **(E)** infections in A549 cells; and upon SARS-CoV-2 high-MOI infection **(F)**, and COVID-19 clinical samples **(G)**. Grey, all genes; Blue non-significant genes (P-value>0.05); red, significant genes (P-value<0.05). The black line is the regression line for all genes. Note that in this case, to allow presenting the maximum number of genes that belong to the tested groups, data filtering, as described in the Methods, was not conducted.

**Figure S11:**
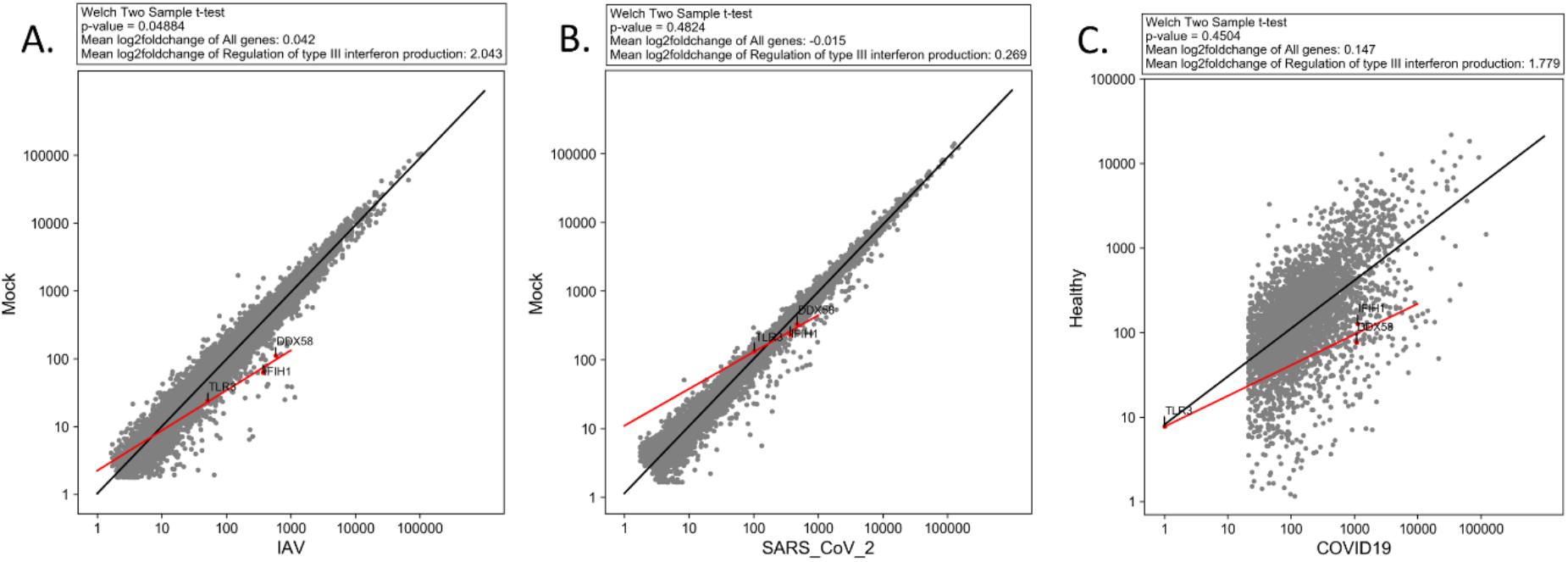
Genes involved in regulation of type III interferon production significantly differentially expressed upon IAV, but not upon SARS-CoV-2, in NHBE cells and COVID-19 patients. Genes involved in the regulation of type III interferon production (red) are significantly upregulated, compared to all genes (grey) upon infection with IAV **(A)**. The test does not reach statistical significance for SARS-CoV-2 infection in A549 cells **(B)** or clinical samples **(C)**. Log scale plots presenting average normalized gene counts. The black and red lines are the regression lines for all genes and regulatory IFN-III genes, respectively. The P-values were obtained using a Welch two-sample T-test. For A, B, and C transcripts with coefficient variation>1 were excluded for the test and presentation. In C, since TLR3 was among the excluded transcripts (with coefficient variation>1), but on the other hand, it is one of the main tested factors, TLR3 was retrieved after filtering.

**Figure S12:**
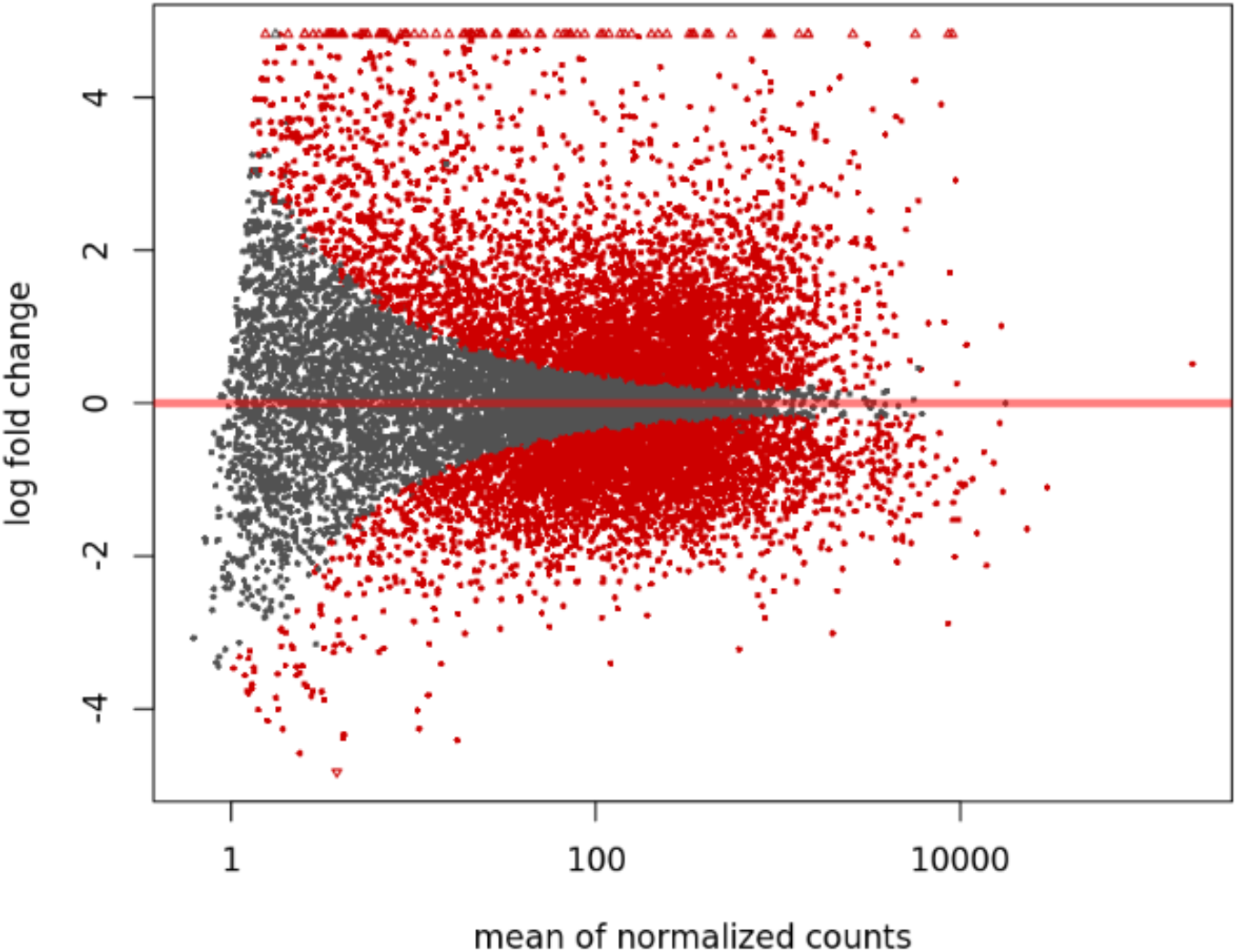
An example for plotMA. The plot shows the log2FoldChange values of SARS-CoV-2 versus mock, plotted against the mean of normalized counts. Data is from GEO GSE147507, series 16.

### Supplementary Tables

Supplementary Table 1: Exported table from BiSEK, showing the normalized counts and DESeq2 results using the default setting, for all samples

Supplementary Table 2: Exported table from BiSEK, showing the normalized counts and DESeq2 results using the default setting, when excluding the outlier sample

Supplementary Table 3: Gene ontology enrichment of significant genes (P-adjusted<0.05), based on Supplementary Table 2

Supplementary Table 4: Exported table from BiSEK, showing the normalized counts and DESeq2 results of SARS-CoV-2 versus Mock (series 1), using the default setting

Supplementary Table 5: Exported table from BiSEK, showing the normalized counts and DESeq2 results of IAV versus Mock (series 9), using the default setting

Supplementary Table 6: Exported table from BiSEK, showing the normalized counts and DESeq2 results of SARS-CoV-2 versus Mock (series 16) using lfcThreshold=1

Supplementary Table 7: Exported table from BiSEK, showing the normalized counts and DESeq2 results of RSV versus Mock (without the outlier mock) (series 8) using lfcThreshold=1

Supplementary Table 8: Exported table from BiSEK, showing the normalized counts and DESeq2 results of HIPV3 versus Mock (without the outlier mock) (series 8) using lfcThreshold=1

Supplementary Table 9: Gene ontology enrichment of significant genes (P-adjusted<0.05), based on Supplementary Table 9

Supplementary Table 10: Gene ontology enrichment of significant genes (P-adjusted<0.05), based on Supplementary Table 10

Supplementary Table 11: Gene ontology enrichment of significant genes (P-adjusted<0.05), based on Supplementary Table 11

